# Development of a Novel Class of Self-Assembling dsRNA Cancer Therapeutics: a Proof of Concept Investigation

**DOI:** 10.1101/2020.04.22.055905

**Authors:** Vishwaratn Asthana, Brett S. Stern, Yuqi Tang, Pallavi Bugga, Ang Li, Adam Ferguson, Anantratn Asthana, Gang Bao, Rebekah A. Drezek

## Abstract

Cancer has proven to be an extremely difficult challenge to treat. Several fundamental issues currently underlie cancer treatment including differentiating self from non-self, functional coupling of the recognition and therapeutic components of various therapies, and the propensity of cancerous cells to develop resistance to common treatment modalities via evolutionary pressure. Given these limitations, there is an increasing need to develop an all-encompassing therapeutic that can uniquely target malignant cells, decouple recognition from treatment, and overcome evolutionarily driven cancer resistance. We describe herein, a new class of programmable self-assembling dsRNA-based cancer therapeutics, that uniquely targets aberrant genetic sequences, and in a functionally decoupled manner, undergoes oncogenic RNA activated displacement (ORAD), initiating a therapeutic cascade that induces apoptosis and immune activation. As a proof-of-concept, we show that RNA strands targeting the EWS/Fli1 fusion gene in Ewing Sarcoma cells that are end-blocked with phosphorothioate bonds and additionally sealed with a 2’-U modified DNA protector can be used to induce specific and potent killing of cells containing the target oncogenic sequence, but not wildtype.

## Introduction

Cancer is the second leading cause of death globally, and was responsible for 8.8 million deaths in 2015 according to the World Health Organization (1). Currently, the gold standard of care for cancer is some combination of chemotherapy, hormonal therapy, targeted molecular therapy, radiation, and/or surgical resection. However, each of these approaches are to varying degrees, non-specific, leading to undesirable side effects on healthy tissue (2). In addition, with many of these therapies, especially targeted molecular therapy, the method of recognition and method of efficacy are intricately coupled. As a result, the choice of target affects the efficacy of therapy, often producing suboptimal results (2). A good example of this is antisense small interfering RNA (siRNA) technology. siRNA is highly specific, targeting strands with sequence complementarity to the therapeutic silencing RNA strand; however, siRNA’s mechanism of action involves cleaving and degrading the target strand. It is entirely possible that the unique cancerous sequence being targeted is not essential for driving the cancerous phenotype and so its degradation has limited benefits. One of the last major issues with conventional therapies is that evolutionary pressure often drives cancerous cells to adopt a resistant phenotype leading to refraction/remission (3). Given these limitations, there is a increasing need for a new class of all-encompassing cancer therapeutics that can uniquely target malignant cells, decouple recognition from treatment, and circumvent cancer resistance.

A fundamental difference between malignant cells and normal tissue is the presence of genetic mutations. Unique mutations can be identified during the pathological staging of biopsy samples using mutation panels or Next Generation Sequencing (4, 5). A method of targeting these genetic mutations, possibly multiple at once, represents an ideal form of personalized medicine and would allow for the selective identification of cancerous cells. Here, we describe a new class of RNA-based cancer therapeutics called ORAD (Oncogenic RNA Activated Displacement) that targets mutated cancerous mRNA in a selective and programmable manner based on simple Watson-Crick thermodynamic base-pairing rules.

The ORAD system is composed of a targeting RNA strand and a complementary DNA protector. As depicted in Figure 1, when the RNA/DNA duplex encounters a wildtype strand with insufficient complementarity, the DNA protector fails to release, leading to no response. However, when the targeted cancerous sequence is encountered, the cancerous mRNA is able to dislodge the DNA protector via strand displacement, producing a therapeutic double stranded RNA (dsRNA) product. We found that by end-blocking the targeting RNA with phosphorothioate bonds to prevent non-specific degradation, and modifying the DNA protector with 2’-U residues to render the RNA/DNA duplex inert prior to opening, premature activation of the ORAD system in cells not harboring the target oncogenic sequence can be prevented.

**Figure 1.**
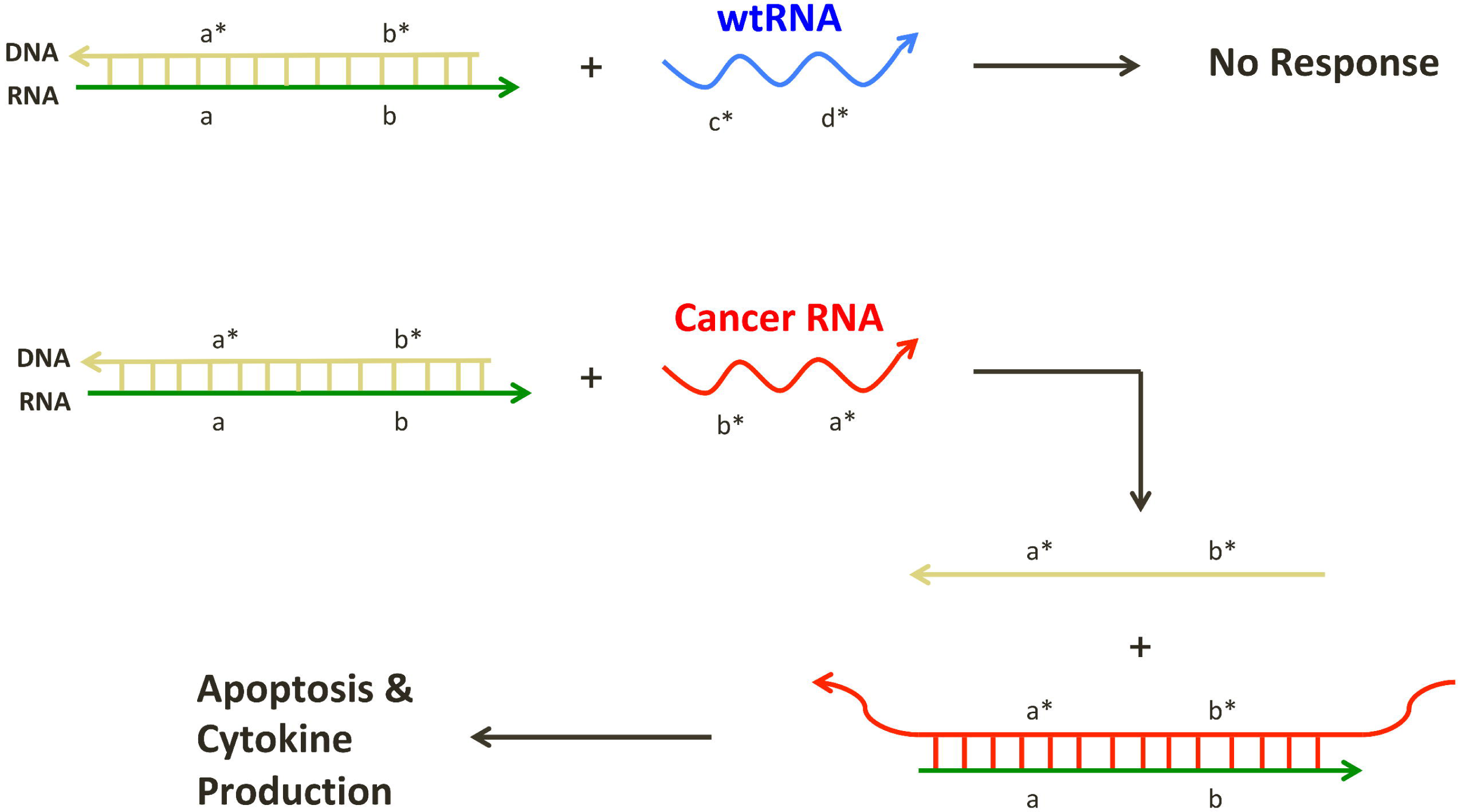
Oncogenic RNA activated displacement (ORAD) schematic. The ORAD system is composed of a targeting RNA strand and a complementary DNA protector. When the RNA/DNA duplex encounters a wildtype strand with insufficient complementarity, the DNA protector fails to release, leading to no response. However, when the targeted cancerous sequence is encountered, the cancerous mRNA is able to dislodge the DNA protector via strand displacement, producing a therapeutic dsRNA product leading to apoptosis and cytokine production. Arrowheads signify 3’ ends. Asterisks signify complementarity.

In general, RNA/RNA base pairing is more thermodynamically favorable than RNA/DNA base pairing, enabling DNA protector displacement in the presence of a cancerous mRNA target even though their sequences are almost entirely homologous (6-10). The use of a DNA protector to confer selectivity was first demonstrated by Zhang et al., 2012 and has been validated to ensure near-optimal specificity across the diverse concentrations, sequence compositions, and salinities that may be encountered intracellularly (11). An in depth explanation of the probe-protector system can be found in the Supplementary Text.

Long dsRNA, or more precisely strands greater than 30 base pairs (bp), are considered foreign elements in eukaryotic cells (12). dsRNA is typically found among viruses possessing dsRNA genomes or dsRNA intermediates during replication. Accordingly, higher-level organisms have adopted mechanisms of identifying long dsRNA and responding to the infectious source (13). In humans, several key proteins recognize long dsRNA. These include protein kinase R (PKR), Toll-like receptor 3 (TLR3), Melanoma differentiation-associated protein 5 (MDA5), retinoic acid-inducible gene I (RIG-I), and 2’-5’-oligoadenylate synthase (OAS). PKR is an intracellular protein that binds dsRNA in a length dependent fashion and induces apoptosis in the host cell to prevent viral propagation (12). TLR3, MDA5, and RIG-I also recognize dsRNA and play an active role in immune activation. TLR3 is a surface receptor expressed primarily on antigen presenting cells, while MDA5 and RIG-I are cytoplasmic helicase receptors expressed in almost all cell types. These three proteins function in innate immunity by recognizing dsRNA and activating NF-kB or interferon regulatory factors, leading to production of inflammatory cytokines, such as type I interferons (14-21). RIG-I and MDA5 also possess caspase recruitment domains (CARD) capable of inducing apoptosis when activated (22). Lastly, OAS, in response to dsRNA, produces 2’-5’-oligoadenylates, which activate ribonuclease L (RNase L) leading to the destruction of both viral and endogenous mRNA in the cell (23, 24).

By producing a long dsRNA product, the ORAD system falsely alerts the cell, and possibly whole body, of a potential viral infection via the aforementioned dsRNA-sensing pathways. Activation leads not only to apoptosis of the target cancer cell but also stimulation of the immune system (via activation of NF-kB or interferon regulatory factors) and subsequent production of inflammatory cytokines. Apoptosis and immune activation represent two independent therapeutic pathways induced by distinct yet slightly overlapping dsRNA-sensing pathways providing a potential means to subvert evolutionarily-driven cancer resistance. In addition, by producing a long dsRNA in the presence of a unique cancer marker, the ORAD scheme functionally decouples recognition from therapy by eliciting a therapeutic affect that is independent of the cancer marker being targeted. This permits the targeting of virtually any uniquely transcribed cancer mRNA with a known sequence.

As a starting cancer model, we have chosen Ewings Sarcoma, an extremely malignant tumor of the bone and soft tissue with an extensively studied fusion gene. Sequences with closer homology to their wildtype counterpart, such as small nucleotide polymorphisms (SNPs), are more difficult to distinguish using ORAD due to the marginal thermodynamic difference between the intended target and wildtype. Cancerous fusion genes and their functional transcripts however, contain a very distinct nucleotide sequence around the fusion site, representing an ideal starting point to test ORAD due to the large thermodynamic difference between the intended target and wildtype. Approximately 90% of Ewing sarcoma cases contain a t(11;22)(q24;q12) chromosomal translocation resulting in the fusion of the EWS gene on chromosome 22 with the FLI1 gene on chromosome 11 (25). The EWS/FLI1 fusion gene produces a functional mRNA transcript that is ultimately translated into the EWS/FLI1 oncogenic fusion protein. Proof of concept tests on the A-673 human Ewing Sarcoma cell line, which expresses the EWS/Fli1 fusion transcript, and corresponding WPMY-1 wildtype cells using ORAD, reveal the potency and selectivity of the system and its potential as an all-encompassing cancer therapeutic (26).

## Methods

### RNA Synthesis

RNA were transcribed using the HiScribe T7 high yield RNA synthesis kit (New England Biolabs) according to manufacturer’s instructions. To prevent the formation of aberrant dsRNA products during T7 RNA transcription, the concentration of MgCl_2_ was limited to 6 mM (27). 100 ng of DNA gBlock (Integrated DNA Technologies) containing the T7 promoter was used as template and transcribed for 48 hours at 37°C. The post-transcription reaction mixture was incubated with 10 units of DNase I (New England Biolabs) at 37°C for 1 hour to remove the gblock template then purified using a RNA spin column (Zymo Research). Purified RNA was then treated with 200 units of calf intestinal phosphatase (New England Biolabs) at 37°C for 24 hours to ensure complete removal of any 5’-triphosphate moieties then purified again using a RNA spin column (Zymo Research). Lastly, RNA yields were determined using a Qubit 3.0 fluorometer (Invitrogen).

For modified RNA synthesis, including 2-thiouridine (s2U), 4-thiouridine (s4U), GU wobble, and 5-methylcytidine (5-mCTP), the natural base was replaced with its modified counterpart at equimolar concentrations and synthesized as described above. Targeting RNA strands end-blocked with phosphorothioate bonds were chemically synthesized by Integrated DNA Technologies (IDT). 2’-fluoro (2’-F) modified RNA was synthesized using the DuraScribe T7 transcription kit (Lucigen) as described in the manufacturer’s protocol. A list of all sequences and primers used can be found in Supplementary Table 1.

### DNA Synthesis and RNA Protection Protocol

DNA was synthesized using standard Taq polymerase (New England Biolabs). For 2’-U scrambled DNA synthesis, a reverse primer only approach was utilized: 5 uL 10x Taq buffer, 1 uL each of 10 mM dNTP (or 2’-U), 1 uL of 100 uM reverse primer only, 2 uL of 50 mM MgCl_2_, 1 uL of 10 ng/uL DNA template, 1 uL of Taq DNA polymerase, and 36 uL of H_2_O (50 uL total). The reaction mixture was then run through the following PCR protocol on the T100 Thermal Cycler (Bio-Rad): 1) denature at 95°C for 3 min, 2) denature at 95°C for 45 seconds, anneal at 55°C for 30 seconds, extend at 72°C for 1 min 30 seconds, 5) repeat steps 2 - 4 30 times, 6) final extension at 72°C for 10 min. Samples were then purified using a DNA spin column (Zymo Research) and quantified using a NanoDrop apparatus (ThermoFisher Scientific). DNA protector strands containing 2’-O methyl modifications were chemically synthesized by Integrated DNA Technologies (IDT).

For high yield synthesis of 2’-U EWS/Fli1 DNA, the following modified Taq protocol was utilized with both forward and reverse primer: 5 uL 10x Taq buffer, 1 uL each of 10 mM dNTP (or 2’-U), 1 uL of 100 uM 5’ monophosphate-modified forward primer, 1 uL of 100 uM 5’ phosphorothioate-modified reverse primer, 2 uL of 50 mM MgCl_2_, 1 uL of 1 ng/uL DNA template, 1 uL of Taq DNA polymerase, and 35 uL of H_2_O (50 uL total). The reaction mixture was then run through the same thermocycling protocol listed above. Synthesized product was subsequently purified using a DNA spin column then digested using lambda exonuclease (New England Biolabs): 10 uL 10x lambda exonuclease buffer, 10 uL lambda exonuclease (50 units), 12 uL dsDNA template, and 68 uL H_2_O (100 uL total) at 37°C overnight. Lambda exonuclease digestion is required to isolate the desired antisense DNA protector from a dsDNA PCR product. Because lambda exonuclease preferentially digests DNA with a 5’-monophosphate, the forward primer, designed to elongate the non-desired sense DNA strand, is modified with a 5’-monophosphate instead of 5’-hydroxyl. In addition, the reverse primer, designed to elongate the desired anti-sense DNA strand, is modified with six phosphorothioate bonds on the 5’ end to inhibit exonuclease digestion (28). Following lambda exonuclease treatment, strands were purified using a DNA spin column then quantified using a NanoDrop apparatus.

To duplex and protect the targeting RNA strands, 0.15 ug/uL of DNA protector and 0.1 ug/uL of targeting RNA (1:1.5 ratio) were thermally annealed in 1x PBS using the T100 thermal cycler based on the following protocol: samples were initially heated to 95°C for 5 minutes, then uniformly cooled to 20°C over the course of 1 hour. For experiments testing and characterizing the 2’-U protected end-blocked targeting RNA (Figure 5), 0.1 ug/uL of RNA was annealed with 0.1 ug/uL of DNA (1:1 ratio) to remove any excess, non-duplexed DNA protector.

**Figure 2.**
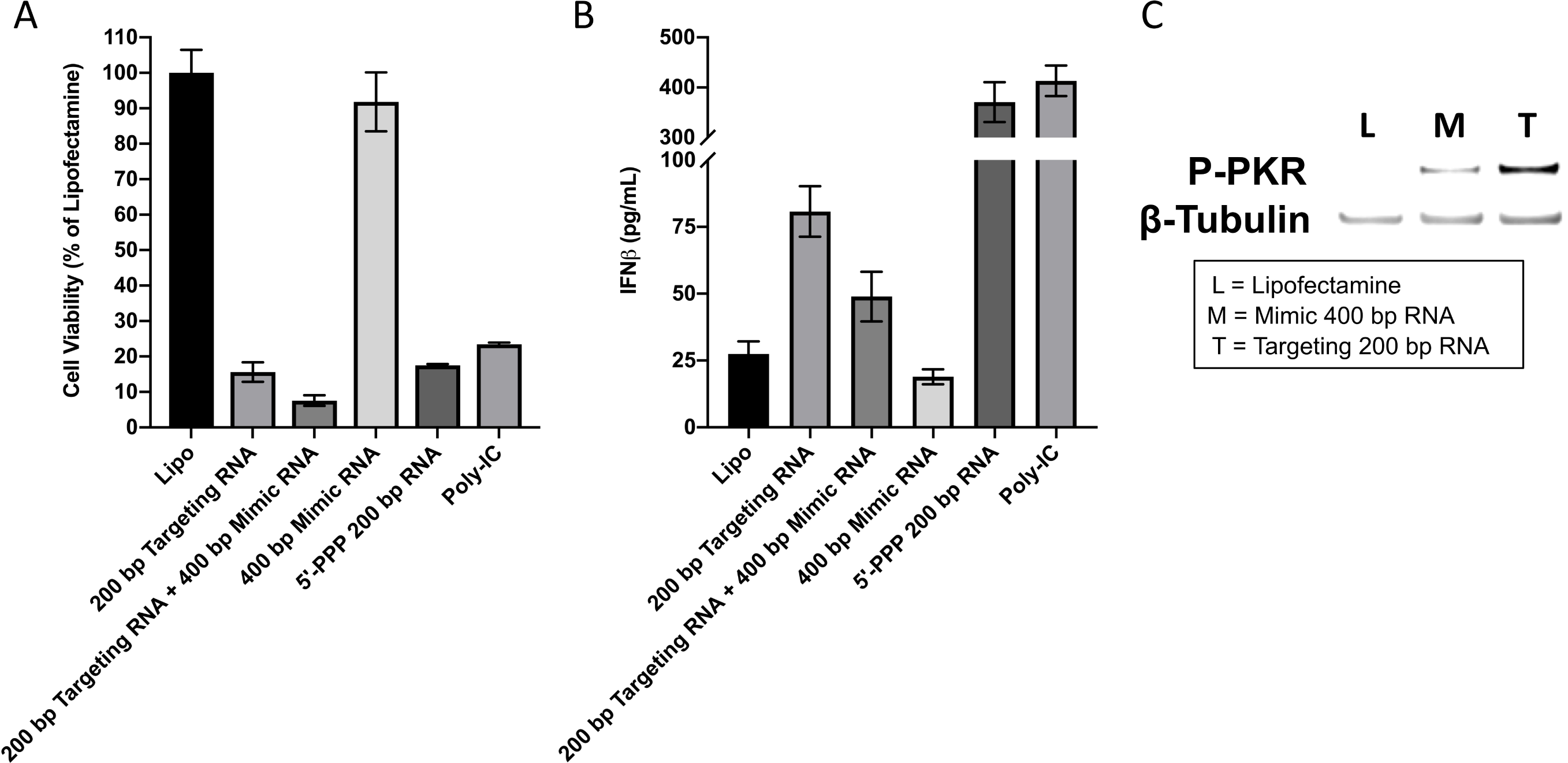
Assessing the cytotoxic and immunogenic potential of the ORAD system. [A] Cytotoxicity and [B] IFNβ cytokine induction levels of 200 bp EWS/Fli1 targeting RNA were compared with levels generated by a reference 400 bp EWS/Fli1 sense RNA strand as well as two established positive controls: poly(I:C) and 5’-triphosphate in A-673 cells. [C] Activation and subsequent phosphorylation of the dsRNA-sensing protein PKR was measured in the presence of lipofectamine (L), 400 bp EWS/Fli1 mimic (i.e. sense) RNA (M), or 200 bp EWS/Fli1 targeting RNA (T), using western blot. Error bars represent the standard deviation of replicate conditions.

**Figure 3.**
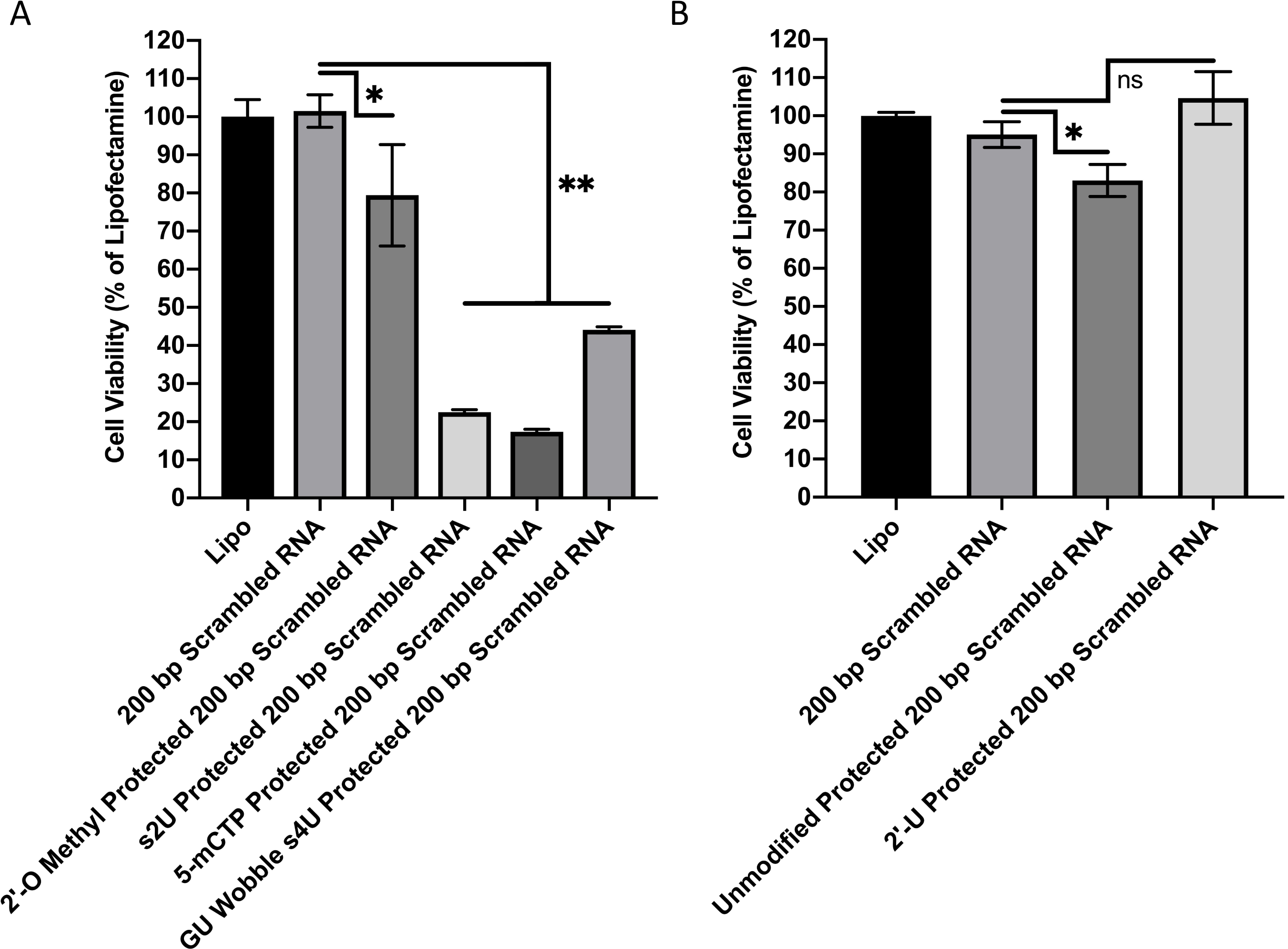
Chemically modifying protector complexes to prevent non-specific cytotoxicity. Scrambled 200 bp targeting RNA strands were protected with either modified [A] RNA or [B] DNA protectors and transfected into A-673 cells using lipofectamine. Only scrambled targeting RNA protected with 2’-U modified DNA were rendered inert and non-toxic. s2U = 2-thiouridine, 5-mCTP = 5-methylcytidine, s4U = 4-thiouridine, 2’-U = 2’-deoxyuridine. Error bars represent the standard deviation of replicate conditions (* = p < 0.05, ** = p < 0.005, ns = non-significant).

**Figure 4.**
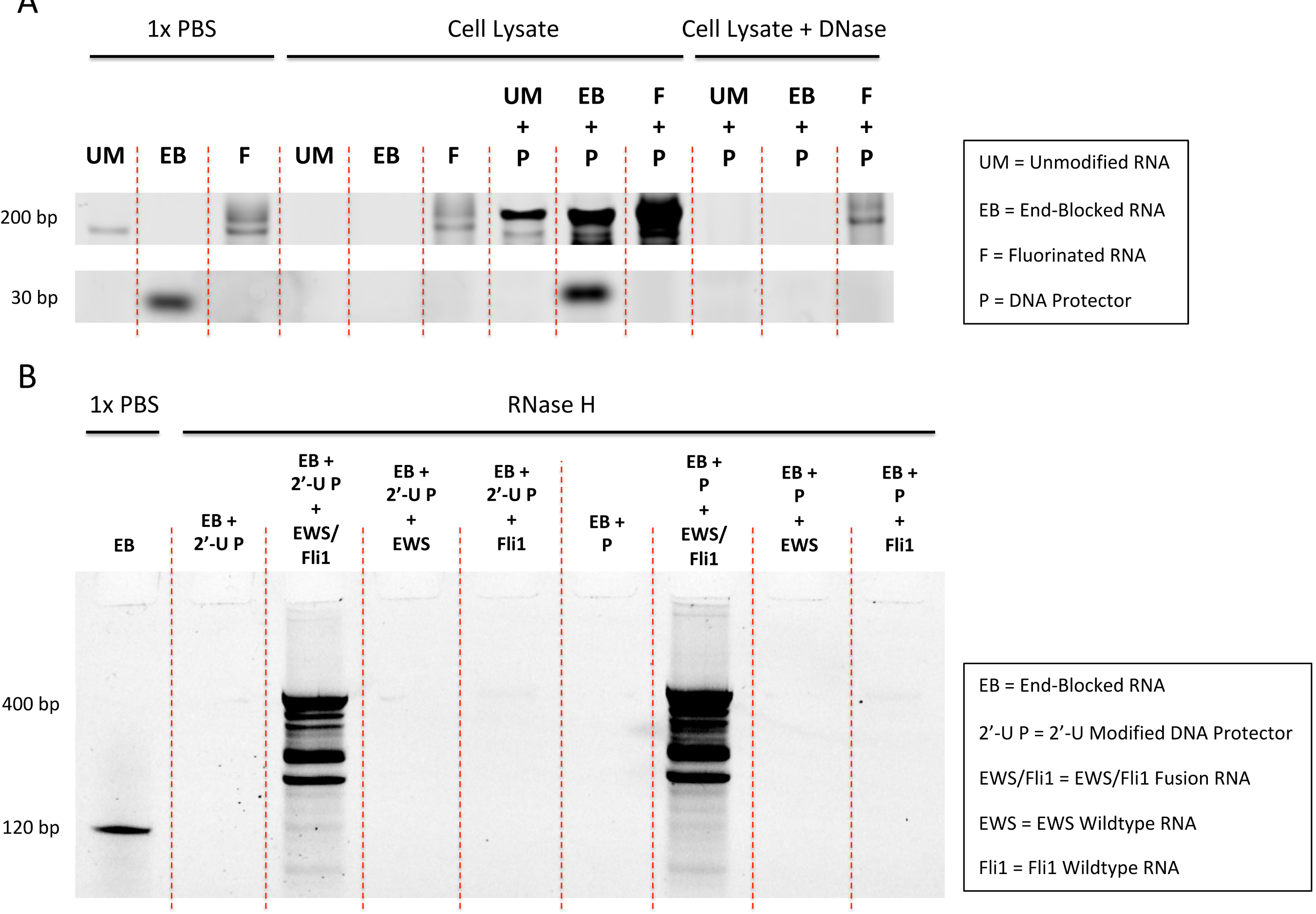
Modifying targeting RNA to resist endo/exonucleolytic degradation and assessing selective displacement extracellularly. [A] 200 bp unmodified (UM), 30 bp chemically synthesized test size end-blocked (EB), and 200 bp 2’-fluorinated (F) EWS/Fli1 targeting RNA were incubated alone or protected with DNA (P) in either 1x PBS or WPMY-1 cell lysate for 48 hours. Post-incubation, select samples were digested with DNase to remove overlapping signal from the DNA protector. All conditions were then run on a denaturing PAGE-Urea gel and visualized using SYBR Gold. The presence of a DNA protector prevents degradation of end-blocked targeting RNA but not unmodified targeting RNA, while 2’-fluorination protects targeting RNA even in the absence of a DNA protector. It should be noted that the chemically synthesized end-blocked RNA, already shortened due to synthetic length restraints, is no longer visible post-DNase digestion because the ends of the end-blocked RNA contain DNA, which when degraded in the presence of DNase, causes the RNA band to shift downwards and off the gel. [B] After establishing end-blocked targeting RNA as a viable candidate, 120 bp 2’-U (2’-U P) and unmodified (P) DNA protected end-blocked and fluorescently-labeled EWS/Fli1 targeting RNA were incubated with 400 bp copies of EWS/Fli1, EWS, and Fli1 mRNA for 48 hours in 1x PBS. The displaced strands were then treated with RNase H and subsequently run on a denaturing PAGE-Urea gel. RNase H should only degrade targeting RNA that is not displaced by the 400 bp RNA transcripts.

**Figure 5.**
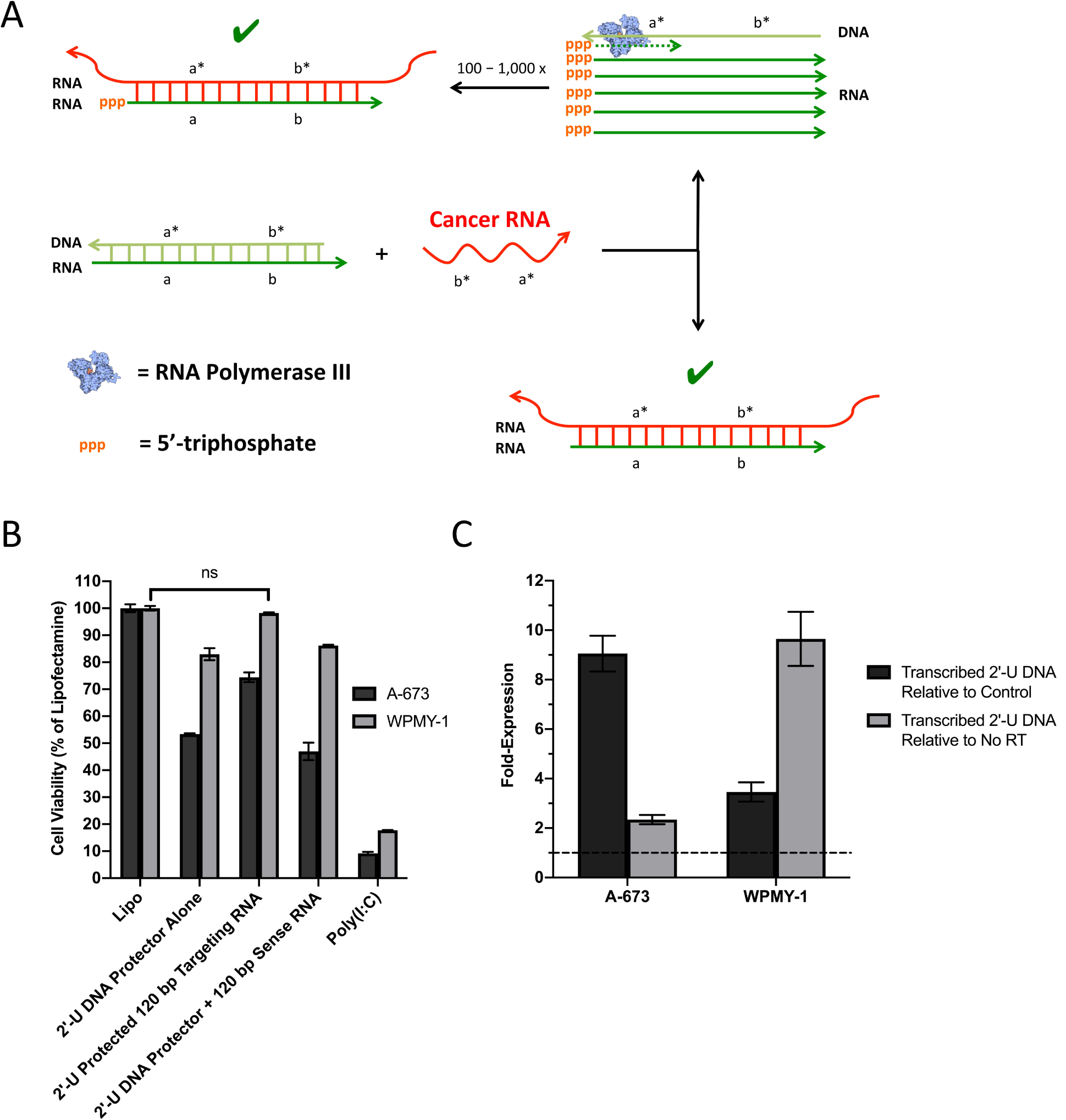
Conferring specificity using 2’-U DNA protected EWS/Fli1 targeting RNA. [A] Hypothesized mechanism of targeting RNA and 2’-U DNA protector induced cytotoxicity. In target cancer cells, cancerous mRNA is able to break the protector seal of the ORAD system and bind to the targeting RNA producing a dsRNA product that activates all the established dsRNA pathways. By binding to the targeting RNA however, the 2’-U DNA protector is released making it a potential substrate for RNA polymerase III. The 2’-U DNA protector then serves as a template for the transcription of hundreds to thousands of potential RNA transcripts that are similar in sequence to the targeting RNA, ultimately amplifying the cytotoxic potential of the ORAD system. The presence of 5’-triphosphate on the transcribed RNA only serves to enhance the effect. [B] 2’-U DNA protected 120 bp end-blocked EWS/Fli1 targeting RNA strands complexed at a 1:1 ratio by mass were transfected into A-673 and WPMY-1 cells using lipofectamine. When complexed at a 1:1 ratio, cell viability in WPMY-1 approaches non-toxic levels with cytotoxicity still apparent in A-673 cells. The use of a non-complementary EWS/Fli1 sense RNA strand with the sense 2’-U DNA protector confers no protective benefit in WPMY-1 cells (ns = non-significant). [C] RT-qPCR was used to quantify transcribed RNA expression levels of EWS/Fli1 2’-U DNA protector in treated cells. In both A-673 and WPMY-1 cells, a weak but significant (greater than unity which is indicated by the dotted line) transcribed 2’-U DNA signal is apparent that exceeds both non-specific signal from control untreated cells as well as signal that originates from DNA, the latter of which was determined by running a no reverse transcriptase (RT) control. Error bars represent the standard deviation of replicate conditions. Arrowheads signify 3’ ends. Asterisks signify complementarity.

### Cell Viability & Cytokine Studies

A-673 and WPMY-1 cells were obtained from ATCC (American Type Culture Collection; Manassas, Virginia) and maintained in Dulbecco’s Modified Eagle Medium (DMEM) supplemented with 10% Fetal Bovine Serum (FBS) and 1% penicillin/streptomycin. Both cells were grown at 37°C in 5% CO_2_. For plating, cells were trypsinized from their flasks and quantified manually using a bright-line hemocytometer (Sigma-Aldrich; St. Louis, Missouri). Replicates of each dilution were then plated on either a 48- or 24-well Corning Costar flat bottom cell culture plate (Thermo Fisher Scientific; Waltham, Massachusetts) at either 1 × 10_4_ cells/well or 5 × 10_5_ cells/well respectively and grown overnight. Cells were then transfected with nucleic acid (RNA and/or DNA) using Lipofectamine RNAiMax (Life Technologies) at a ratio of 0.3 ug nucleic acid / 1 uL Lipofectamine. The total amount of nucleic acid added to each experimental well was 3.15 ug of RNA and/or DNA / 50,000 cells, requiring approximately 10.5 uL of Lipofectamine RNAiMax to effectively deliver this dose. Experiments testing the chemically synthesized 120 bp targeting RNA required double the nucleic acid / Lipofectamine dose to induce an appropriate response (a detailed discussion of ORAD dosing can be found in the Supplementary Text). After a 48 hour incubation, cells were stained with 4′,6-diamidino-2-phenylindole (DAPI) and quantified using microscopic cytometry as described previously (29). For cytokine studies, supernatant was extracted from the cell plates just prior to cell staining then processed and measured using a human IFN-β ELISA kit (PBL Assay Science).

For extended transfection studies, cells were plated onto 24-well cell culture plates at 5 × 10_5_ cells/well then allowed to adhere overnight. Cells were transfected as described above then given 48 hours to incubate after which the media was replaced with fresh media with one phosphate buffered saline (PBS) wash in between. Cells were transfected once again and given another 48 hours to incubate after which cell viability was assessed.

### Western Blot

A-673 cells were plated onto a 6-well cell culture plate at 5 × 10_6_ cells/well then allowed to adhere overnight. Cells were transfected as described above then given 24 hours to incubate. Following incubation, cells were lysed with RIPA buffer (Santa Cruz) as described in the manufacturer’s instructions then centrifuged at 10,000g for 10 minutes to remove cell debris. Protein concentrations were calculated using a bicinchoninic acid (BCA) protein assay (Pierce) according to the manufacturer’s instructions. Cell lysates (20 ug) were diluted in 4x LDS buffer (Life Technologies) with 5% Beta-mercaptoethanol. Samples were denatured by heating to 95°C for 5 minutes and cooled to room temperature. Proteins were resolved by SDS-PAGE on 4-12% gradient gels (Invitrogen) using MOPS running buffer (Life Technologies), and transferred to polyvinylidene fluoride (PVDF) membranes. Membranes were blocked for one hour at 40 RPM on shaking platform with a 2:1 ratio of Odyssey Blocking Solution (Li-Cor) to PBS with 0.05% Tween-20 (PBS-T). Anti-pPKR T446 (Abcam, ab32036) and anti-β tubulin (Developmental Studies Hybridoma Bank, E7) primary antibodies were diluted 1:1000 in a solution of PBS-T with 0.1% Bovine Serum Albumin (BSA). Primary antibodies were then detected using goat anti-rabbit 680 nm (Rockland Immunochemical, RL6111440020.5) and goat anti-mouse 800 nm secondary antibodies (Rockland Immunochemical, RL6111450020.5) respectively after incubating in PBS-T + 0.1% BSA for two hours. Membranes were imaged on an Odyssey Classic Imager (Li-Cor).

### Cell Lysate Extraction

Cultured cells were brought into suspension using standard cell culture protocol, centrifuged at 250g for 5 minutes to form a cell pellet, then washed once with PBS. Cells were subsequently resuspended in cell lysis buffer (920 µL H_2_O, 50 µL 1M Tris-HCl pH 7.4, 10 µL 10% sodium dodecyl sulfate, 10 µL Igepal CA-630, 8.77 mg NaCl, 5 mg sodium deoxycholate, and 10 µL protease inhibitor cocktail) at a ratio of 1 mL cell lysis buffer per 10_6_ cells then incubated for 15 minutes on an orbital shaker. Lysed cells were centrifuged at 12,000g for 10 minutes. The remaining supernatant is the cell lysate.

### RNA Extraction and RT-qPCR

Cell lysate was extracted, after incubating cells for 24 hours in condition, using the cell lysate extraction protocol listed above. Cell lysate was purified using a RNA spin column then treated with uracil-DNA glycosylase (5 µL UDG, 5 µL of 10x UDG reaction buffer, 25 uL purified cell lysate, and 15 uL of H_2_O) (New England Biolabs) for 1 hour at 37°C followed by DNase I (10 µL DNase I, 10 µL of 10x DNase I buffer, 50 uL UDG-treated cell lysate, and 30 uL of H_2_O) for 1 hour at 37°C. Following UDG and DNase I incubation, samples were purified again using an RNA spin column.

After digesting 2’-U and genomic DNA, RNA levels were quantified using either a conventional 1-step RT-qPCR or a modified 2-step RT-qPCR protocol, both using the iTaq universal SYBR green one-step kit (BioRad). For 1-step RT-qPCR, the following reaction components were mixed together: 5 uL of 2x iTaq universal SYBR green reaction mix, 0.125 uL iScript reverse transcriptase, 0.5 uL of 10 uM forward and reverse primer, 1 uL cell lysate, and 2.875 uL H_2_O. The RT-qPCR samples were then measured using the CFX96 Touch Real-Time PCR Detection System (BioRad) using the following thermocycling protocol: 1) reverse transcribe at 50°C for 10 minutes, 2) denature DNA and activate Taq polymerase at 95°C for 1 minute, 3) denature DNA at 95°C for 10 seconds, 4) anneal and extend at 62°C for 20 seconds followed by fluorescence capture, and 5) repeat steps 3 – 4 35 times.

### Assessing RNA Stability in Cell Lysate and RNase H Displacement Assay

For assessing RNA stability in cell lysate, 2.5 uL of 0.1 ug/uL protected or unprotected targeting RNA was incubated in 2.5 uL of cell lysate at 37°C for 48 hours. The digested samples were then mixed 1:1 (v/v) with formamide (Sigma-Aldrich) and run on a denaturing PAGE-Urea gel (ThermoFisher Scientific) in 1x TBE buffer at 60°C. For RNase H displacement assays, 1 uL of 0.05 ug/uL DNA protected fluorescently-labeled targeting RNA was incubated with 1 uL of 0.1 ug/uL 400 bp EWS or Fli1 RNA in 1x PBS and incubated at 37°C for 48 hours. Following incubation, the samples were digested using RNase H in the following reaction mixture at 37°C for 20 minutes: 2 uL sample, 0.5 uL 10x RNase H reaction buffer, 0.5 uL RNase H, and 2 uL H_2_0. The digested samples were then mixed 1:1 (v/v) with formamide and run on a denaturing PAGE-Urea gel in 1x TBE buffer at 60°C.

### Statistics

All statistical analyses were performed using GraphPad Prism 8 Software. Significance was set at α = 0.05. Comparisons between groups were assessed using one-way ANOVA.

## Results

### Determining the Cytotoxic and Immunogenic Potential of the ORAD System

It has been shown that dsRNA-binding proteins, like PKR, respond synergistically to multimonomer binding, with longer strands likely producing a stronger response (12). Preliminary tests suggest that *in vitro* transcribed (Fig S1) EWS/Fli1 antisense targeting RNA strands 200 bp in size, with 100 bp of the targeting RNA strand complementary to the EWS portion of the EWS/Fli1 fusion mRNA and the other 100 bp complementary to the Fli1 portion, were the most potent out of a range of targeting RNA varying in length from 20 bp to 390 bp (Fig S2). These 200 bp EWS/Fli1 targeting RNA were delivered into A-673 cells in the absence of a DNA protector to assess their cytotoxic potential (Fig 2a). They were also tested against a reference *in vitro* transcribed 400 bp RNA sense strand, which is a truncated mimic of the EWS/Fli1 mRNA transcript, as well as two established positive controls: 5’-triphosphate and poly(I:C). The 200 bp targeting RNA is shown to be extremely potent, inducing a greater than 80% reduction in cell viability, while the reference 400 bp RNA sense strand is non-toxic. At this targeting RNA length, we can expect PKR, RIG-I, and OAS to be active but not MDA5, which tends to activate in the presence of significantly longer dsRNA, typically kilobases or larger (30-32).

IFNβ production was then assessed using supernatant from the treated wells (Fig 2b). IFNβ is a type I interferon that serves an important role in cancer immunotherapy and is a relatively sensitive marker of dsRNA pathway activation. In general, IFNβ induction levels closely mirror cytotoxic trends. To see if the ∼10-20% of cells that survived initial treatment were resistant to the ORAD system, we treated A-673 cells either once or twice with therapeutic RNA spaced by the appropriate incubation period (Fig S3). After 48 hours, A-673 cell viability had been reduced by ∼90%, however a repeat administration and additional 48 hour incubation reduces cell viability by another 95%, representing a greater than two-order of magnitude drop in overall cell viability. These results not only indicate the extreme potency of the ORAD system, but also imply that those cells that survive initial treatment have not developed resistance to the therapeutic RNA. To confirm dsRNA-specific pathway activation, PKR autophosphorylation was measured via western blot revealing a robust and specific induction pattern in the presence of 200 bp EWS/Fli1 targeting RNA versus 400 bp EWS/Fli1 mimic (i.e. sense) RNA (Fig 2c). Altogether, data measuring cell viability, IFNβ expression, and PKR activation suggest that the generation of a long dsRNA product is indeed producing the robust response being observed.

Having validated the cytotoxic potential of the ORAD system, we ran strand displacement simulations of the 200 bp EWS/Fli1 targeting RNA now sealed with a DNA protector. Secondary structure analysis revealed that initial variants of the 200 bp targeting RNA/DNA, with the fusion-site located directly in the middle of the strand (100 bp complementary to EWS and Fli1 each), were forming trimeric states with homologous wildtype sequences. To prevent this, the targeting fusion site was shifted towards the 3’ end of the targeting RNA in order to kinetically lock the strands (Supplementary Text).

### Modifying Protector Seal to Prevent Non-Specific Cytotoxicity

Initial tests of the newly designed and synthesized EWS/Fli1 fusion-shifted targeting RNA strands in a wildtype WPMY-1 cell line demonstrated non-specific cytotoxicity when a DNA protector was utilized at a 1:1.5 targeting RNA to DNA protector ratio (data not shown). We hypothesized that the RNA/DNA hybrids of the ORAD system might function as a pathogen-associated molecular pattern (PAMP), and as a result induce non-specific immune activation. Accordingly, various modifications were tested on the protected RNA duplexes with the goal of rendering them fully inert. While modifying the targeting RNA could interfere with the therapeutic pathway of the ORAD system by altering binding to dsRNA-sensing proteins, adapting the protector seal would be far less restrictive. Thus, several modified deoxynucleotides, ribonucleotides, and altered Watson-Crick base pairs that could potentially replace canonical bases and base pairs in the protector were compared to determine which modifications could help the duplex evade detection by PAMP-receptors. These modified bases and altered base pairs include 2-thiouridine (s2U), 4-thiouridine (s4U), GU wobble, 5-methylcytidine (5-mCTP), 2’-deoxyuridine (2’-U), and 2’-O methyl (33-35).

To avoid potential confounding effects from the EWS/Fli1 targeting RNA, we synthesized a new 200 bp scrambled RNA strand, as well as corresponding modified RNA or DNA protectors, to isolate the protective effects of each modification. 2-thiouridine-, 5-methylcytidine-, and GU wobble 4-thiouridine-containing scrambled RNA protectors were synthesized using conventional RNA transcription, while 2’-U-containing scrambled DNA protectors were synthesized using a modified Taq PCR protocol. 2’-O methyl-containing DNA protectors cannot be enzymatically synthesized using PCR so strands modified with ∼10% 2’-O methyl-containing GTPs were chemically synthesized instead.

The newly synthesized scrambled duplexed strands were subsequently transfected into A-673 cells. We found that neither 2’-O methyl, s2U, 5-mCTP, or GU wobble with the s4U modification was sufficient to shut down non-specific cytotoxicity (Fig 3a). However, 2’-U modified DNA proved to be almost completely inert (Fig 3b), making it the ideal protector modification moving forward for the RNA/DNA scheme.

EWS/Fli1 targeting RNA, now protected with 2’-U modified DNA, were again tested in WPMY-1 control cells, however preliminary results indicated that the DNA seal failed to confer any protective benefit (data not shown).

### Modifying Targeting RNA to Resist Degradation

Two possibilities were considered regarding the failure of the new 2’-U protected targeting RNA: either the targeting RNA was degrading in the cell cytoplasm, which would break the protector’s kinetic lock, leading to premature activation of the ORAD system, or the inclusion of the 2’-U moiety altered base-pairing thermodynamics in a way that ultimately reduced the strength of the protector seal.

We first focused on inhibiting RNA degradation in the cytoplasm, via endo- and exo-nucleases, while minimizing alterations to the targeting RNA that could potentially reduce therapeutic efficacy (36, 37). Two candidate modifications were found that could potentially be used to inhibit RNA degradation: 2’-fluorination and phosphorothioate-backbone incorporation (28, 38). Because endonucleases canonically cut at purine base residues, 2’-fluorine modifications at these sites should inhibit not only exonucleases but also endonucleases, even in the absence of a protector (38). We hypothesized that end-blocking RNA with phosphorothioate (PS) bonds would inhibit exonucleases, while adding a DNA protector would protect against endonucleases, in the latter case, by forming a double-stranded nucleic acid complex (28). The addition of DNA bases to the end of the targeting RNA strand should accentuate any protective benefit the PS bonds may confer. 2’-fluorinated RNA was synthesized using the Y639F mutant T7 RNA polymerase, which is able to incorporate non-canonical bases like 2’-F (39). PS-bonds cannot be incorporated using conventional transcription, so end-blocked PS strands were chemically synthesized.

To test the resistance of the modified strands to degradation, 200 bp unmodified RNA, 200 bp fluorinated RNA, or chemically synthesized, shortened (30 bp test size), end-blocked RNA were incubated in WPMY-1 cell lysate for 48 hours. After incubation, strands were run on a denaturing PAGE-Urea gel and stained with SYBR gold (Fig 4a). The RNA strands alone are stable in 1x PBS over 48 hours, but almost entirely break down in the presence of cell lysate, with the exception of fluorinated RNA. As expected, in its protected state, the end-blocked RNA is also resistant to degradation. DNase was used to remove the overlapping signal from the DNA protector, revealing the fluorinated RNA underneath, and further showing that DNA protection alone is insufficient to protect unmodified RNA in cell lysate. It should be noted that the chemically synthesized, 30 bp test size, end-blocked RNA is no longer visible post-DNase digestion because the ends of the end-blocked RNA contain DNA, which when degraded in the presence of DNase, causes the RNA band to shift downwards and off the gel.

After running preliminary cytotoxicity tests in cells, the 2’-F RNA we synthesized was found to be too nonspecifically toxic (data not shown) and was not pursued further, leaving the end-blocked RNA as the primary candidate moving forward. By design, the end-blocked RNA contains two to three PS bonds at either end, along with four to five overlapping DNA bases. Because PS-modified RNA strands need to be chemically synthesized using phosphoramidite solid phase synthetic processes, they can be made no longer than 120 bp with current technology (40).

Having addressed the degradation issue, we next sought to verify that the 2’-U protected end-blocked targeting RNA still followed thermodynamic simulations and bound only to the target EWS/Fli1 mRNA cancerous sequence and not the EWS or Fli1 mRNA wildtype sequences. This was accomplished using an RNase H assay to gauge selective displacement (see Fig S4 for a schematic). RNase H is an endonuclease that cleaves the RNA strand in an RNA/DNA duplex. Because the DNA protector seal is supposed to remain bound to the targeting RNA in the presence of both wildtype sequences, the RNA/DNA duplex remains a substrate for RNase H, leading to targeting RNA degradation. However, in the presence of the target cancerous sequence, the DNA protector is displaced leading to the formation of a dsRNA complex, which is not an adequate substrate for RNase H. Accordingly, the targeting RNA remains intact and available for subsequent detection.

The protected end-blocked RNA were incubated in the presence of either the desired EWS/Fli1 target sequence or the EWS or Fli1 wildtype sequences for 48 hours in 1x PBS then treated with RNase H (Fig 4b). Only EWS/Fli1 RNA is capable of displacing the DNA seal, indicated by resistance to RNase H digestion and corresponding preservation of fluorescent signal. EWS and Fli1 wildtype are incapable of removing the DNA protector, leading to RNase H digestion and loss of signal. No difference in protector performance can be seen between the 2’-U and unmodified DNA protectors signifying that 2’-U modified bases do not have significantly altered base-pairing thermodynamics. Altogether, this suggests that RNA degradation in the cell cytoplasm, and breaking of the protector’s kinetic lock, likely led to premature activation of the 2’-U DNA protected targeting RNA strands in the control WPMY-1 cells, and that this process can be inhibited by end-blocking the targeting RNA.

### Testing and Characterizing 2’-U Protected End-Blocked Targeting RNA

Observational data of early tests utilizing the 2’-U EWS/Fli1 DNA protector were surprising, in that the EWS/Fli1 2’-U DNA protector alone was found to induce cytotoxicity, which was not the case with the scrambled 2’-U DNA protector tested earlier. We realized that if 2’-U DNA mediated cytotoxicity was sequence specific, it is theoretically possible that the sense 2’-U DNA protector is getting transcribed in the cytoplasm into antisense targeting RNA, via an RNA polymerase. Recently, RNA polymerase III, which is typically thought to reside in the nucleus, was found localized in the cytoplasm functioning as a DNA sensor (41). When displaced from the RNA/DNA duplex by the target cancer mRNA sequence, the 2’-U DNA protector is potentially made available for transcription by RNA polymerase III leading to linear amplification of the targeting RNA strand and increased potency of the ORAD system (Fig 5a).

With the 2’-U DNA protector potentially more cytotoxic than the targeting RNA strand but partially uncomplexed as part of the initial RNA/DNA duplex ratio of 1:1.5, an adjustment was made lowering the duplex ratio down to 1:1. Through these changes, the DNA protector of the ORAD system demonstrated intracellular specificity for the first time (Fig 5b). As expected, the 2’-U DNA protector alone was toxic in the A-673 and WPMY-1 cells, but when duplexed at a 1:1 ratio with targeting RNA, cell viability in WPMY-1 returned to near 100% with cytotoxicity still apparent in the A-673 cells. Perhaps most convincing is the complexing of the 2’-U DNA protector with a non-complementary sense RNA strand. The sense RNA strand cannot seal the 2’-U DNA protector, causing the cytotoxicity profile for this condition to closely mirror the 2’-U DNA protector alone condition. Supernatant from the treated wells were also extracted and tested for IFN-β production. When the targeting strands are protected, IFN-β is potently induced in the A-673 target cell line versus the control WPMY-1 cells (Fig S5). It should be noted that the cytotoxic potential of the ORAD system appears to have been reduced after shortening the size of the targeting RNA to 120 bp from 200 bp as well as end-blocking the RNA with both DNA bases and phosphorothioate bonds.

The previously reported data strongly supports the RNA polymerase III hypothesis, though not directly. In an attempt to mechanistically prove RNA polymerase III activity, we sought to detect and quantify the transcribed RNA product that the 2’-U DNA protector would generate in the presence of RNA polymerase III using RT-qPCR. In both A-673 and WPMY-1 cells, a weak but significant (greater than unity) transcribed 2’-U DNA signal was detected that exceeds both non-specific signal from control untreated cells as well as signal that originates from DNA, the latter of which was determined by running a no reverse transcriptase control (Fig 5c). In target A-673 cells, this signal was approximately 9-fold higher than the corresponding control condition, and in WPMY-1 this signal was approximately 3.5-fold higher.

## Discussion

The proof of concept ORAD system described herein represents a potentially new class of cancer therapeutics based on the principle of self-assembling dsRNA. The system is a potent inducer of both cytotoxicity and cytokine production and can selectively target cancerous cells that express unique fusion genes while sparing normal tissue. By producing a long dsRNA product in the presence of a unique cancer marker, the ORAD scheme functionally decouples recognition from therapy by eliciting a therapeutic effect that is independent of the cancer marker being targeted. This permits the targeting of virtually any uniquely transcribed cancer mRNA with a known sequence.

A similar approach has been attempted before using 40 bp antisense RNA strands designed to be complementary to fragments flanking the fusion site of an oncogene (42). When bound to the target oncogene but not wildtype mRNA, a dsRNA product sufficient in length to activate PKR is generated. One of the issues with this approach however is that the length of antisense RNA used must be restricted, which would likely reduce potency. Furthermore, the antisense RNA needs to be constitutively expressed at high levels using a transfected plasmid (42). Another method using RNA hairpin displacement has been attempted, but met with limited success (43). RNA hairpins are generally less stable and consistent than the protected probe scheme listed here, often leading to increased nonspecific activation and off-target effects (11). Lastly, use of preformed dsRNA or poly(I:C), especially as an immune adjuvant in clinical trials, has become increasingly popular. However, targeting and specificity require the expression of unique extracellular antigens that homing vectors can target (44, 45). It is possible that either these unique antigens are not be available for targeting, are also expressed on healthy cells, or are downregulated as a form of evolutionary resistance, limiting the potential use of this form of therapy. By virtue of its design, the ORAD system overcomes many of the aforementioned issues.

We show that *in vitro* transcribed targeting RNA strands 200 bp in length can reduce cell viability by up to 90% in cells that express a target cancer fusion mRNA sequence while an equivalent dose of unprotected 400 bp EWS/Fli1 mimic (sense) RNA, intended to serve as a negative control, is minimally toxic. This suggests that the potent response induced by the 200 bp EWS/Fli1 is in part governed by sequence specificity and not the result of off-target binding. In addition, the 200 bp ORAD targeting RNA was more cytotoxic than both 5’-triphosphate RNA and poly(I:C) which are strong inducers of RIG-I and PKR (14, 46). Supernatant extracted from treated wells show that IFNβ induction levels closely mirror cytotoxic trends with the exception of pre-complexed 200 bp EWS/Fli1 targeting RNA + 400 bp mimic RNA, which produced lower levels of IFNβ than expected. This is likely the result of the overwhelming cytotoxicity of the pre-complexed dsRNA hindering the production of IFNβ. Selective PKR activation in the presence of 200 bp targeting RNA was observed via detection of phosphorylated PKR protein, while 400 bp mimic RNA showed only slight non-specific activation, most likely via the formation of minimal, non-contiguous intramolecular dsRNA segments generated via secondary hairpin formation. Unfortunately, a robust method to detect other endogenously activated dsRNA-sensing proteins including RIG-I and OAS3 could not be established in the current experimental framework. Altogether, the data indicate that the formation of a dsRNA product is indeed responsible for producing the pronounced response that is being observed.

Literature suggests that in the 100 – 200 bp size range, we can expect several dsRNA-sensing pathways, including PKR, RIG-I, OAS, and Dicer, to activate but not MDA5 (47). It is possible that targeted cytotoxicity may be even more pronounced at significantly longer targeting RNA lengths by recruiting additional dsRNA-sensing proteins. By activating multiple, somewhat redundant dsRNA-sensing pathways, that induce two distinct therapeutic pathways (apoptosis and immune activation), the ORAD system is potentially resilient to evolutionary resistance acquired either upstream or downstream of the recognition portion of the dsRNA signaling transduction cascade. Because cancerous mutations, especially driver mutations, are signatures of the cancerous phenotype and not easily downregulated, the ORAD system’s recognition of unique cancer mRNA represents another means to prevent evolutionarily-driven cancer resistance (48). The ability to target multiple mutations at once only serves to strengthen this effect. This phenomenon is confirmed to an extent with the repeat transfection test that was conducted on cancer target cells that survived initial treatment. The initial 10% of cells that “evaded” ORAD-induced cytotoxicity were further reduced in number by approximately an order of magnitude upon repeat administration likely indicating that those cells that survive initial treatment have not developed resistance to the therapeutic RNA. With multiple dosing regimens, cytotoxicity could theoretically reach 100%.

In general, assessment of both cytotoxicity and cytokine production proved to be the most sensitive assays for testing the efficacy of the ORAD system during each stage of the development process. A more robust assay to detect therapeutic activation, including western blot or use of a fluorescent reporter cell line, is not available to assess the multifaceted components of the ORAD system. Western blot, for example, is not sufficiently quantitative to detect the subtle intracellular response to each ORAD modification, while a fluorescent reporter cell line cannot take into the account the multiple dsRNA-sensing pathways that converge to induce apoptosis and immune activation. Altogether, these alternative assays are too narrow in focus to assess the cumulative response of each ORAD-induced pathway.

After demonstrating the success of long dsRNA in inducing cytotoxicity and activating the innate immune system, thus confirming the *therapeutic* arm of the ORAD system, we focused on engineering DNA protectors that would confer specificity to the targeting RNA strands in order to validate the *diagnostic* arm of the ORAD system. To maintain selectivity and prevent trimer formation, the ORAD RNA/DNA hybrids were modified so that the targeting RNA fusion-site was shifted towards the 3’ end of the strand, kinetically locking the system. For specificity studies, the WPMY-1 cell line was chosen as a control comparator given its origin from histologically normal tissue (albeit immortalized) and closer resemblance to actual wildtype cells encountered in the human body (49). This cell line was chosen in place of siRNA or gene knockdown of EWS/Fli1 mRNA in A-673 cells, as these methods significantly alter the transcriptome and proliferative potential of the cell, likely because the EWS/Fli1 fusion protein is in part responsible for driving the cancerous phenotype in Ewing Sarcoma (50).

After running cytotoxicity studies of the newly designed and synthesized EWS/Fli1 fusion-shifted targeting RNA strands in control WPMY-1 cells, the DNA protected targeting RNA complexes were found to be non-specifically cytotoxic. We realized that the RNA/DNA hybrids of the ORAD system might function as a PAMP. In response to the very long cytoplasmic RNA/DNA duplexes that were exogenously being introduced into the cell, inflammasomes, including the NLRP3 inflammasome, might be triggering and inducing non-specific cytotoxicity. The NLRP3 inflammasome has been shown to recognize RNA/DNA hybrids of bacterial origin that have gained access to the cytoplasm. Upon recognition and activation, NLRP3 inflammasomes induce the production of IL-1β, maturation of IL-18, and stimulation of a form of inflammatory cell death known as pyroptosis (51, 52). To render the RNA/DNA duplexes inert, various modifications to the protector were tested, including the incorporation of 2-thiouridine (s2U), 4-thiouridine (s4U), GU wobble, 5-methylcytidine (5-mCTP), 2’-deoxyuridine (2’-U), and 2’-O methyl. Only insertion of 2’-U modifications into the DNA protector rendered scrambled RNA/DNA duplexes non-toxic in A-673 cells. 2’-deoxyuridine is unique in that it contains structural components from both RNA and DNA. While 2’-U has a DNA sugar lacking a 2’-hydroxyl, it contains an RNA uracil base. Though induction and subsequent resolution of PAMP activation was not directly assessed, 2’-U may create a hybrid structure that is not an adequate substrate for pathogen-detection systems in the cell including not only the NLRP3 inflammasome but also traditional dsRNA-sensing pathways. Future studies investigating the specific components of the PAMP activation pathway, such as the NLRP3 inflammasome, will help elucidate their various contributions to cell viability and immunogenicity in the setting of the ORAD system. Ultimately, the ability of 2’-U to render the ORAD complexes inert makes it the ideal candidate moving forward for the RNA/DNA scheme (53).

After testing fusion-shifted EWS/Fli1 targeting RNA protected with newly synthesized 2’-U modified DNA in WPMY-1 cells, we found that the DNA seal failed to confer any protective benefit. We speculated that either the targeting RNA was degrading in the cell cytoplasm, which would break the protector’s kinetic lock, or that inclusion of the 2’-U moiety was reducing the strength of the DNA protector seal. We focused first on inhibiting RNA degradation using either 2’-fluorination or phosphorothioate-backbone incorporation. After synthesizing the appropriate strands, unmodified, fluorinated and end-blocked phosphorothioate modified RNA were incubated in cell lysate for 48 hours and run on a denaturing PAGE-Urea gel. Unmodified RNA was susceptible to degradation regardless of whether it was duplexed with a DNA protector while 2’-F was resilient to degradation with or without a protector seal. The end-blocked phosphorothioate modified RNA was only resistant to degradation when duplexed to the DNA protector confirming the dominant role of exonucleases in degrading and potentially prematurely activating strands of the ORAD system.

Because 2’-F RNA was found to be too nonspecifically toxic, we were left with end-blocked, PS-modified RNA as the primary candidate for protecting the targeting RNA of the ORAD system. By design, the end-blocked RNA contains two to three PS bonds at either end, along with four to five overlapping DNA bases. Phosphorothioate bonds substitute a sulfur atom for a non-bridging oxygen in the phosphate backbone of an oligonucleotide strand. This renders the inter-nucleotide linkage resistant to nuclease degradation. However, the number of PS bonds that can be incorporated into a strand is limited due to nonspecific toxicity of the bonds (28). An additional limitation is that these strands need to be chemically synthesized limiting the size of the targeting RNA to no more than 120 bp.

Selective displacement of the newly synthesized 2’-U DNA protected, 120 bp fusion-shifted, end-blocked targeting RNA was then tested using an RNase H assay. Incubating the strands in 1x PBS for 48 hours with truncated *in vitro* transcribed copies of the target EWS/Fli1 and wildtype EWS and Fli1 sequences confirms the specificity of the ORAD strands for the intended target, indicated by resistance to RNase H digestion and corresponding preservation of fluorescent signal with the EWS/Fli1 target sequence and loss of signal due to RNase H digestion in the presence of the EWS and Fli1 wildtype sequences. The complex band pattern observed between 120 bp and 400 bp in the presence of EWS/Fli1 is likely due to abnormal migration of the thermodynamically stable but partially double-stranded RNA complex (120 bp is duplexed fully while 280 bp is single-stranded). An extended discussion regarding the RNase H displacement assay can be found in the Supplementary Text.

Having modified the ORAD strands to prevent premature activation in the cell cytoplasm, we tested various combinations of the newly modified, end-blocked 120 bp synthetic EWS/Fli1 targeting RNA with a complementary 2’-U DNA protector to assess any effects alterations to the RNA would have on cytotoxicity. Interestingly, we noticed that the EWS/Fli1 2’-U DNA protector alone was inducing cytotoxicity, which was not the case with the scrambled 2’-U DNA protector used in Figure 3. We hypothesized that an additional enzyme—RNA polymerase III—may be contributing to the toxicity of the 2’-U DNA by transcribing it in the cytoplasm into what is effectively targeting RNA. RNA polymerase III is typically thought to reside in the nucleus where it transcribes rRNA, tRNA, and other small RNAs (54). However, recent literature suggests that RNA polymerase III is also localized in the cytoplasm where it functions as a DNA sensor, transcribing foreign DNA into RNA with a 5’-triphosphate, which can subsequently be detected by RIG-I. In particular, sequences rich in adenine and thymine base residues were found to be adequate substrates for RNA polymerase III in the cytoplasm (41). Though it’s not clear how, 2’-U may function similarly.

Based on the proposed interaction between RNA polymerase III and the ORAD system depicted in Figure 5a, both the 2’-U DNA and targeting RNA can be viewed as primary therapeutic components with each serving as the protector for the other. Accordingly, the targeting RNA to DNA protector ratio was brought down from 1:1.5 (DNA protector in 0.5x excess) to 1:1 (DNA protector present in equimolar amounts). For the original ratio, which had the 2’-U DNA protector in 50% excess of the targeting RNA, a small but non-trivial percent of 2’-U DNA was left un-complexed making it available for potential transcription.

With the 1:1 RNA to DNA ratio adjustment, we observed the resolution of non-specific cytotoxicity in WPMY-1 cells suggesting that the RNA and DNA strands of the ORAD system were now appropriately sealed and inert in control cells. The sequence specific effect of the protector complex was confirmed using a non-complementary sense RNA strand in conjunction with the 2’-U DNA protector, which together was inadequate to seal the duplex and prevent cytotoxicity. It should be noted that the cytotoxicity of the 2’-U protected targeting RNA is slightly reduced in A-673 cells relative to the 2’-U DNA protector alone when it should theoretically remain unchanged. We suspect that the displacement reaction is not 100% complete in the target cell line and that further tuning of the toehold regions may be required. In addition, the reduced length of the targeting RNA as well end-blocking with both PS bonds and DNA bases appear to have notably reduced overall cytotoxicity. While reducing the length of the targeting RNA can theoretically reduce the cytotoxic potential of the ORAD system by decreasing the number of dsRNA-binding monomers that can attach to the final targeting RNA complex, it is not clear why end-blocking the strands is also inhibitory. It is possible that certain dsRNA-sensing proteins are incapable of accessing the ORAD targeting RNA when end-blocked. Though crystal structures of PKR suggest that the protein is capable of binding dsRNA internally, helicases like RIG-I may need to initiate binding at the 5’ or 3’ ends of the strand (55-58). Although end-modification incorporation is currently necessary to prevent RNA degradation and chemical synthesis is required to insert these modifications thereby limiting the size of the targeting RNA, it may be possible in the near future to chemically synthesize longer strands, incorporate end-blocking modifications via *in vitro* transcription, or utilize a different set of end-blocking modifications that minimally alter the targeting RNA strand to preserve cytotoxicity at levels seen in Fig 2a, while simultaneously maintaining specificity.

In an attempt to mechanistically prove the RNA polymerase III mechanism, RT-qPCR was used to detect and quantify the transcribed RNA product that the 2’-U DNA protector would generate in the presence of RNA polymerase III. After running total RNA extraction on A-673 and WPMY-1 cells treated with and without EWS/Fli1 2’-U DNA protector, incubating the extract with Uracil-DNA Glycosylase to remove 2’-U DNA as well as DNase to remove genomic DNA, and running gene specific primer RT-qPCR, we detected a trace (greater than unity) but unique RNA signal in cells treated with the 2’-U DNA protector versus untreated control cells. The transcribed RNA signal was also stronger than any signal originating from DNA, which was determined by running a no reverse transcriptase control. To ensure that the signal enhancement seen in the 2’-U DNA protector condition versus untreated control was not the result of changes in Fli1 expression levels, a modified 2-step RT-qPCR was performed confirming the identity of the amplified product as the RNA transcribed 2’-U DNA (Supplementary Text).

The aforementioned data strongly supports the RNA polymerase III hypothesis, albeit indirectly. If the proposed interaction of the ORAD system with RNA polymerase III is indeed true, it would raise the cytotoxic potential of the system considerably due to the linear amplification of therapeutic agent. In addition, even though RNA polymerase III potentially enhances the cytotoxicity of the system, it is not essential. For cancerous cells that do not express RNA polymerase III or have it knocked down as a form of evolutionary resistance, the primary targeting RNA strand is still present to enact the original design of the system. More studies are required to verify the RNA polymerase III hypothesis or determine possible alternative mechanisms by which 2’-U DNA is transcribed intracellularly. Additional work is also necessary to establish the precise role of chemical modifications and sequence of the DNA protector in the transcription potential of exogenous ORAD DNA in the cytoplasm. Further exploration of the proposed 2’-U mechanism may elucidate novel therapeutic targets or future avenues for cancer treatment.

## Conclusions

ORAD represents a proof of concept system to induce specific and potent killing of cells containing a target oncogenic sequence but not wildtype, decouple recognition from treatment, and overcome evolutionarily driven cancer resistance. With further advances in RNA synthesis methodology including, but not limited to, chemical synthesis of longer strands or ability to incorporate end-blocking modifications via *in vitro* transcription, ORAD treatment with longer 2’-U DNA protected targeting RNA and an appropriate delivery vehicle has the potential to induce a robust response *in vivo* and one day improve progression-free and/or overall survival in the setting of a clinical trial. In theory, the targeting strands of the ORAD system only need to be administered once as immune surveillance and memory can suppress tumor recurrence and metastasis. However if a tumor were to recur, this form of personalized medicine can be repeated as long as a biopsy and genomic sequence is attainable (59). The ability to potentially target any cancer type is also strongly compelling. We hope our self-assembling dsRNA cancer therapeutics will one day improve the survival and quality of life for cancer patients, and introduce a paradigm shift in how we view and treat cancer, by placing a special emphasis on what makes cancer fundamentally unique—genetics.

## Acknowledgements

We would like to thank the Miller lab for providing the Nikon Eclipse Ti-E inverted fluorescent microscope, the Zhang lab for providing feedback on the DNA protected RNA designs of the ORAD system, and the Drezek lab for editorial assistance. We would also like to thank the Baylor College of Medicine Medical Scientist Training Program (MSTP) for providing training support throughout the duration of the project.

