## Supplementary Text for "Development of a Novel Class of Self-Assembling dsRNA Cancer Therapeutics: a Proof of Concept Investigation"

### Supplementary Information

#### Probe-Protector Mechanism and Thermodynamic Simulations of the ORAD System

The ORAD system is composed of a targeting RNA strand and a complementary DNA protector. The use of a DNA protector to confer selectivity was first demonstrated by Zhang et al., 2012 (11). Initial variations of the probe-protector system developed by Zhang et al. involved the hybridization of an approximately 30 bp probe, sealed by a complementary 20-25 bp protector, to an intended target. This process is initiated at the exposed 5' single-stranded region of the probe (also known as a toehold and is typically 5 bp in length), proceeds through branch migration, and is completed upon total dissociation of the protector at the 3' region of the probe. The toeholds allow the displacement reactions to proceed with fast kinetics.

The underlying principle of the probe-protector system is that it is possible to design complementary protector strands that hybridize less favorably to the diagnostic/therapeutic probe strand than the intended target while binding more favorably than a spurious target based on the thermodynamics of Watson-Crick base-pairing. Overall, the system has been validated to ensure near-optimal specificity across the diverse concentrations, sequence compositions, and salinities that may be encountered intracellularly.

The ORAD design is similar to initial variations of the probe-protector system with several fundamental differences. First, the ORAD system targets an RNA strand using an RNA probe and a DNA protector instead of targeting a DNA strand using a DNA probe and DNA protector. In general, RNA/RNA base pairing is more thermodynamically favorable than RNA/DNA base pairing, enabling DNA protector displacement in the presence of a cancerous mRNA target even though their sequences are almost entirely homologous (6-10). Second, while the ORAD system contains a toehold for target binding on the 3' end of the probe, it does not contain a corresponding 5' toehold to allow the protector strand to potentially re-anneal when in equilibrium. Lastly, the strands of the ORAD system are significantly longer (on the order of 200 bp) than the initial variants of the probe-protector system in order to induce potent cytotoxicity when the system is activated upon dsRNA product formation. While the ability to thermodynamically resolve SNPs is effectively lost at this length, the ability to discriminate unique fusion sequences is preserved as elaborated upon below.

In its initial form, the fusion site of the ORAD targeting RNA lied directly in the middle of the strand (100 bp complementary to EWS and Fli1 each). Secondary structure analysis using NUPACK suggests that with this particular design, the protected RNA would displace fully in the presence of the EWS/Fli1 mRNA target. However, due to the large size of the targeting RNA strand, in the presence of the EWS or Fli1 mRNA wildtype sequences, a trimeric state would form where targeting RNA, its DNA protector, and the target mRNA were all bound (Fig S6) (8,60). The formation of the simulated trimeric state would lead to non-specific activation in wildtype cells given that the region of double strandedness forming between the targeting RNA and its complementary wildtype mRNA sequence would exceed the 30 bp detection limit of dsRNA-sensing proteins such as PKR, defeating the purpose of a protector strand.

To prevent trimeric formation, the region complementary to the fusion-site was shifted towards the 3’ end of the targeting RNA. With this new scheme, 180 bp of the targeting RNA was now complementary to Fli1 and 20 bp was complementary to EWS. Now with a complementary 195 bp DNA protector sealing the targeting RNA strand, a 5 bp 5’ non-homologous region designated for protector binding only, and a 5 bp toehold on the 3’ end designated for target binding only, the targeting RNA/DNA hybrid was kinetically locked (Fig S7). In other words, EWS mRNA, which functions as the initiator of protector displacement, cannot proceed past the 20 bp mark unless it is fused to Fli1. Even if EWS were to stably bind the 20 bp region without getting displaced by the DNA protector, the region of double-strandedness is insufficient in length to induce dsRNA-sensing proteins like PKR. While the majority of the targeting RNA is now complementary to Fli1, Fli1 mRNA is kinetically incapable of initiating strand displacement due to the 5 bp non-homologous region on the 5’ end and the 20 bp EWS region on the 3’ end.

It should be noted that while the fusion-shifted RNA/DNA duplexes are kinetically locked, they are not thermodynamically sealed. That is to say, if Fli1 were able to access its complementary region on the targeting RNA strand, the DNA protector would be displaced. Similarly, it is not possible to re-simulate selective displacement of the DNA protector in the presence of the fusion-gene target versus wildtype using currently available nucleic acid hybridization models because they are limited to thermodynamic simulations and not kinetics. The kinetics of the reaction however, can be extrapolated from the literature. In a 2012 paper by Zhang et al., the kinetics of strand displacement in the absence of a toehold region was found to be on the order of months

(11). The kinetics may be even slower with 5' and 3' seals (non-homologous buffer nucleotides). This timeframe is more than sufficient for the system to be considered adequately sealed.

**ORAD Dosing**

Cells were transfected with nucleic acid (RNA and/or DNA) using Lipofectamine RNAiMax (Life Technologies) at a ratio of 0.3 ug nucleic acid / 1 uL Lipofectamine. The ideal ORAD therapeutic dose was found to be approximately 3.15 ug of RNA and/or DNA / 50,000 cells, with approximately 10.5 uL of Lipofectamine RNAiMax required to effectively deliver this dose. We found that the optimum ORAD therapeutic window was narrow and was limited in the upper regime by non-specific Lipofectamine toxicity and in the lower regime by therapeutic RNA efficacy, with potency dropping sharply in response to a marginal decrease in therapeutic RNA dosing (limiting a sufficiently expansive dose-response study). It should be noted however that due to diminished cytotoxicity, double the nucleic acid / Lipofectamine dose was required for experiments testing the chemically synthesized 120 bp targeting RNA in order to induce an appreciable cytotoxic response. Overall, it is likely that a critical threshold of therapeutic RNA per cell is required to induce an all-or-nothing apoptotic response. In the future, newer delivery vehicles may be able to deliver higher concentrations of targeting RNA with minimal transfection-associated cytotoxicity.

**Inhibiting RNA Degradation**

In general, most endonucleases that target RNA cleave ssRNA (36,37). Endonucleases that cleave dsRNA, like RNase III, which is responsible for the formation of siRNA, would only act after the formation of dsRNA and therapeutic activation. Endonucleases that cleave RNA/DNA, like RNase H, which is responsible for cleaving the RNA portion of Okazaki fragments, are localized primarily in the nucleus (61). It should be noted that while anti-sense oligonucleotides (ASO) are known to trigger mRNA degradation in the nucleus via RNase H, there is a belief that ASO-mediated mRNA degradation in the cytoplasm is also attributable to RNase H. It has been found, however, that ASO-mediated degradation of mRNA in the cytoplasm is likely mediated by steric blockage of mRNA translation, followed by transport and degradation in mRNA-degradation bodies instead of via RNase H (62). Altogether, only endonucleases that cleave ssRNA (instead of RNA/DNA or RNA/RNA) would be capable of digesting the targeting RNA strands of the ORAD system via endonucleolytic pathways, however the use of a DNA protector should prevent this. Accordingly, we sought to inhibit RNA exonuclease degradation using either 2'-fluorination or phosphorothioate-backbone incorporation.

**Special Considerations Regarding the RNase H Selective Displacement Assay**

The RNase H test we ran demonstrates that 2'-U incorporation does not significantly alter base-pairing thermodynamics. It also validates the fusion-shifted design and use of a kinetic lock. Had a trimeric complex formed in the presence of wildtype, the strands of the ORAD system would have been partially protected from RNase H leading to incomplete degradation and preservation of trace signal.

It should be noted that while running the extracellular displacement assay in cell lysate instead of 1x PBS would have been more representative of conditions found intracellularly, the exogenous 400 bp RNAs are not resistant to degradation and would eventually breakdown in cell lysate allowing the RNA/DNA strands to re-anneal. In the absence of the exogenous 400 bp RNAs, endogenous levels of the corresponding mRNA strands are too low to visualize on a gel. Similarly, while assessing selective displacement of the ORAD complex intracellularly instead of on a gel (using tools such as molecular beacons) would provide direct evidence of target hybridization, we found that the concentration of target mRNA and resultant number of hybridization events did not generate a sufficiently strong signal to overpower the background bleed through of the fluorophore/quencher pair.

**Verifying the RNA Polymerase III Hypothesis and Validating the Identity of the RNA Transcribed 2'-U DNA RT-qPCR Product**

A trace (greater than unity) but unique RNA signal in cells treated with the 2'-U DNA protector versus untreated control cells was detected in cells using RT-qPCR, presumably via RNA polymerase III. Though expression levels of the transcribed RNA were expected to be much higher, there are two potential reasons as to why this might not have been the case. First, because the start and stop site of RNA polymerase III on the 2'-U DNA protector is unknown, it is possible that the primer set utilized was non-optimal for amplifying the given target. Alternatively, only a small number of transcripts were actually produced by RNA polymerase III using the 2'-U DNA protector as a template but the reason the 2'-U system is so potent in cells is because the transcribed RNA retains its 5'-triphosphate. Regardless, a unique intracellular signal that very likely originated via RNA polymerase III had been detected. Attempts at characterizing transcribed RNA levels, if any, from unmodified DNA instead of 2'-U modified DNA proved unsuccessful as the background DNA signal could not be sufficiently lowered without Uracil-DNA Glycosylase.

To ensure that the signal enhancement seen in the 2'-U DNA protector condition versus untreated control was not the result of changes in Fli1 expression levels (the primer pair overlaps not only with the targeting RNA but also Fli1 mRNA), we modified our RT-qPCR protocol to introduce directionality bias. Because the gene specific primer pair used in the initial RT-qPCR test is incapable of differentiating the antisense targeting RNA from its sense EWS/Fli1 or Fli1 mRNA counterpart, preferential amplification of the desired targeting RNA strand versus its complement is required to determine which strand is contributing primarily to the detected signal. This was achieved using an adapted 2-step RT-qPCR setup. First, reverse transcription was performed on the extracted RNA samples using a gene specific reverse primer only. This would ensure cDNA production primarily of the desired targeting RNA strand and not its complement. The reverse transcriptase was then inactivated using heat at which point a gene specific forward primer was added and standard qPCR was performed. If the primary product in the 1-step RT-qPCR protocol was indeed targeting RNA, then the use of a reverse primer only during cDNA synthesis would lead to a stronger signal than the use of a forward primer only during cDNA synthesis. The opposite would be the case if the primary product in the 1-step RT-qPCR protocol was Fli1 mRNA. The reverse primer only signal was indeed higher than the forward primer only signal in both cell types, indicating not only that the amplified product is the targeting RNA and not Fli1 mRNA, but also that both cell types have a mechanism to transcribe the 2'-U DNA protector into targeting RNA, likely via RNA polymerase III (Fig S8). These findings are supported by NCBI BLAST data, which confirm that no other endogenous sequences are targets of the targeting RNA primer pair. A summary of the protocol and its reaction components is provided below:

"For the modified 2-step RT-qPCR protocol, the following reaction components were mixed together: 5 uL of 2x iTaq universal SYBR green reaction mix, 0.125 uL iScript reverse transcriptase, 0.5 uL of either 10 uM forward or reverse primer only, 1 uL cell lysate, and 2.875 uL $H_2O$. The reaction mixture was then reverse transcribed at 50°C for 10 minutes then heated to 95°C for 5 minutes to inactivate the reverse transcriptase. After inactivation, 0.5 uL of either 10 uM forward or reverse primer (whichever was not included during the initial setup phase) was added to the reaction mixture then run on the real-time qPCR apparatus using the previously described 1-step RT-qPCR protocol, albeit without the reverse transcriptase step."

#### Supplementary Figures

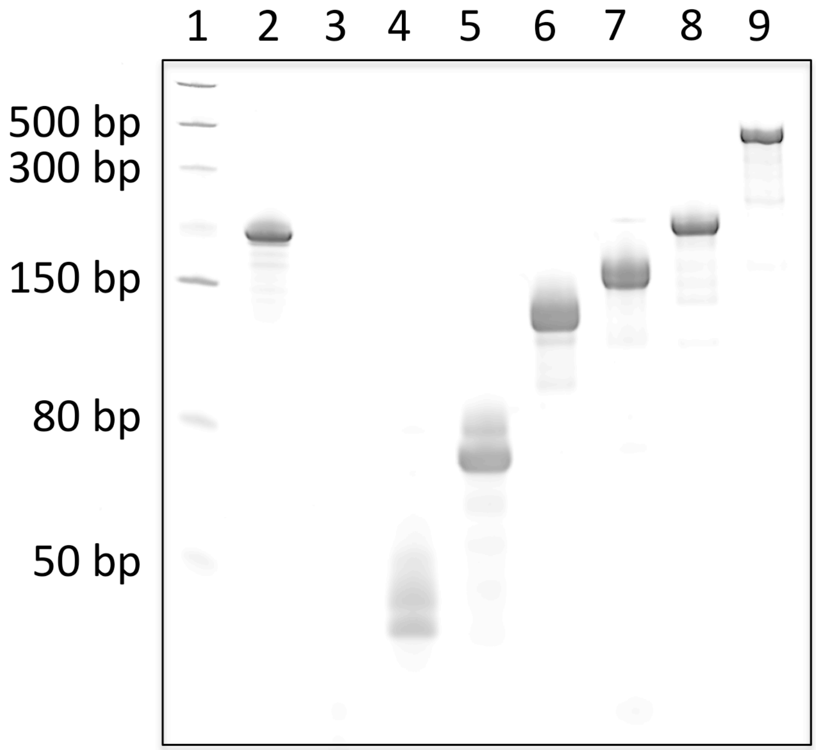


**Figure S1. Transcribing targeting RNA strands of various lengths.** gBlock DNA templates of various lengths were transcribed, DNase treated, spin-column purified, then run on a denaturing PAGE-Urea gel and visualized using SYBR Gold. Lane 1 = ssRNA ladder, lane 2 = 200 bp scrambled RNA, lane 3 = 10/10 EWS/Fli1 targeting RNA, lane 4 = 20/20 EWS/Fli1 targeting RNA, lane 5 = 35/35 EWS/Fli1 targeting RNA, lane 6 = 60/60 EWS/Fli1 targeting RNA, lane 7 = 70/70 EWS/Fli1 targeting RNA, lane 8 = 100/100 EWS/Fli1 targeting RNA, lane 9 = 200/190 EWS/Fli1 targeting RNA. It should be noted that transcript concentrations of 10/10 EWS/Fli1 targeting RNA were low making it difficult to visualize using gel electrophoresis. In addition, the faint smear in each transcribed RNA lane was found to form post-CIP digestion and not immediately post-RNA transcription indicating that the smeared products were the result of non-specific digestion and not spurious RNA transcription.

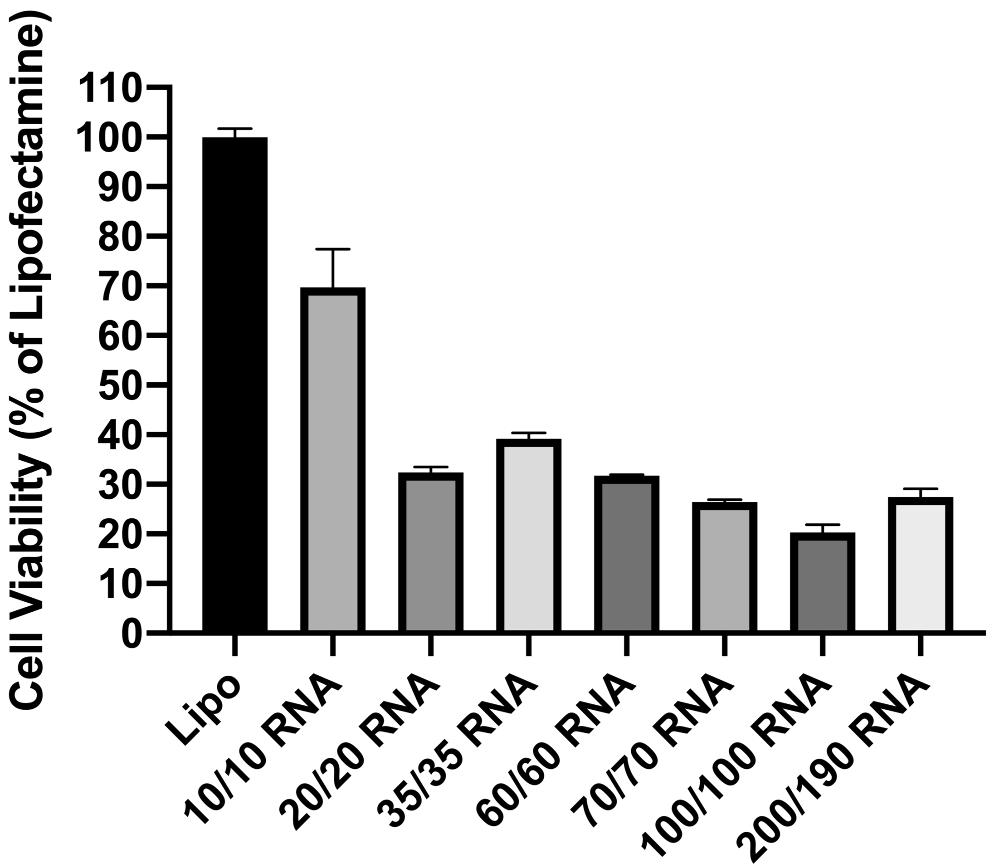


**Figure S2. Determining optimum targeting RNA size of the ORAD system.** Cytotoxicity of EWS/Fli1 targeting RNA strands ranging in length from 20 bp to 390 bp were assessed in A-673 cells in the absence of a DNA protector. Though subtle, potency appears to peak around 200 bp. Error bars represent the standard deviation of replicate conditions.

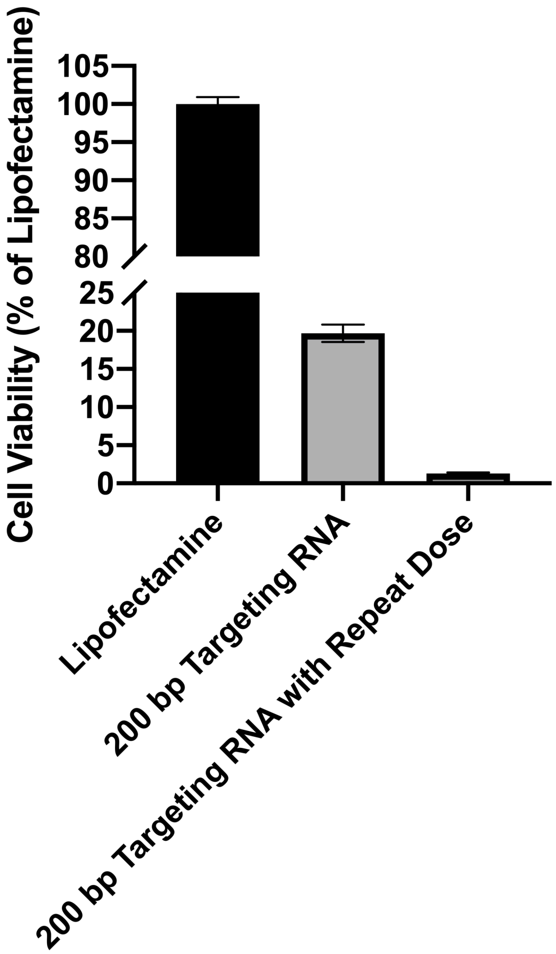


**Figure S3. Assessing resistance to the ORAD system.** Cell viability of A-673 cells treated either once or twice with 200 bp EWS/Fli1 targeting RNA was measured. A-673 cells dosed twice with appropriate incubation spacing did not show a diminished response to the therapeutic indicating that those cells that survive initial treatment have not developed resistance to the therapeutic RNA. Error bars represent the standard deviation of replicate conditions.

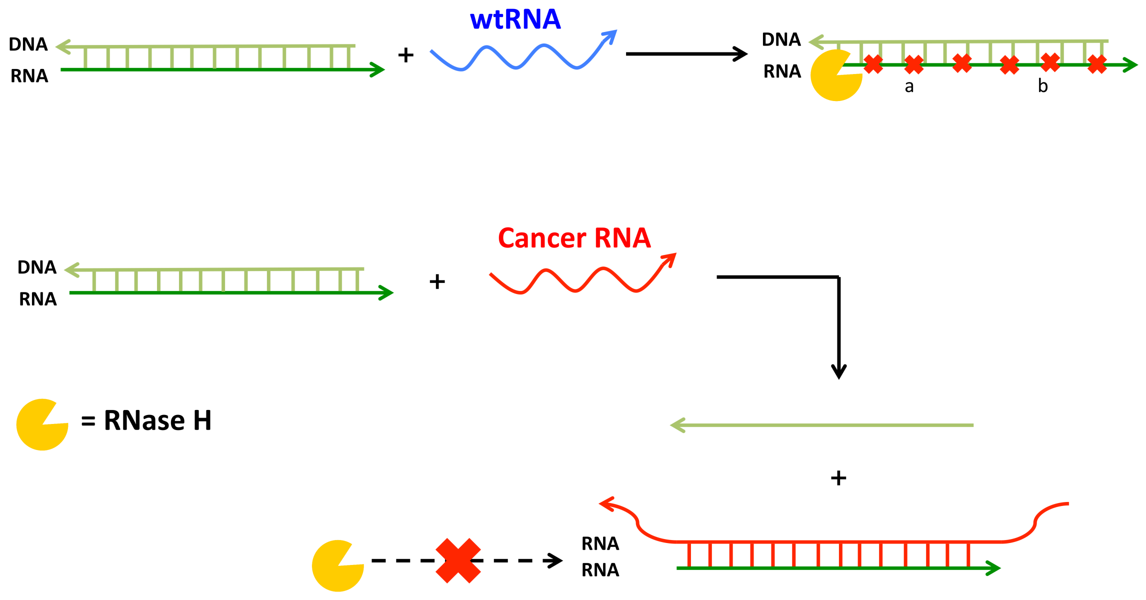


**Figure S4. Schematic overview of RNase H displacement assay.** RNase H is an endonuclease that cleaves the RNA strand in an RNA/DNA duplex. Because the DNA protector seal is supposed to remain bound to the EWS/Fli1 targeting RNA in the presence of both the EWS and Fli1 wildtype sequences, the RNA/DNA duplex remains a substrate for RNase H, leading to targeting RNA degradation. However, in the presence of the EWS/Fli1 target cancerous sequence, the DNA protector is displaced leading to the formation of a dsRNA complex, which is not an adequate substrate for RNase H. Accordingly, the targeting RNA remains intact and available for subsequent detection. Arrowheads signify 3' ends.

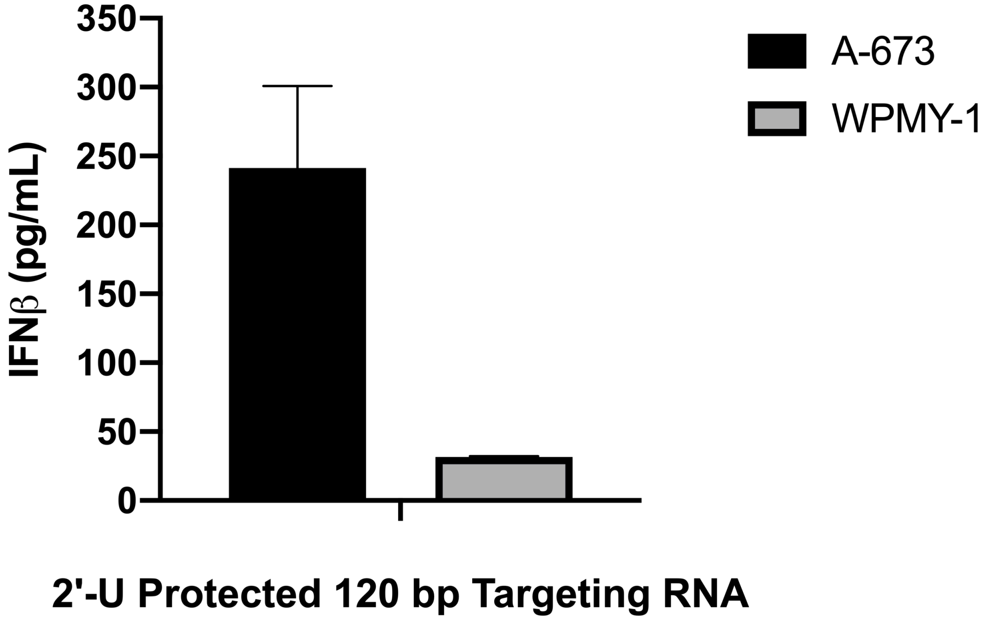


**Figure S5. IFNβ cytokine levels in target A-673 versus control WPMY-1 cells treated with strands of the ORAD system.** Supernatant from A-673 and WPMY-1 cells treated in Figure 5b were extracted and used to quantify IFNβ cytokine levels induced in response to the 2'-U DNA protected 120 bp end-blocked EWS/Fli1 targeting RNA strands of the ORAD system. IFNβ induction is significnatly more pronounced in target A-673 cells versus WPMY-1 control cells. Error bars represent the standard deviation of replicate conditions.

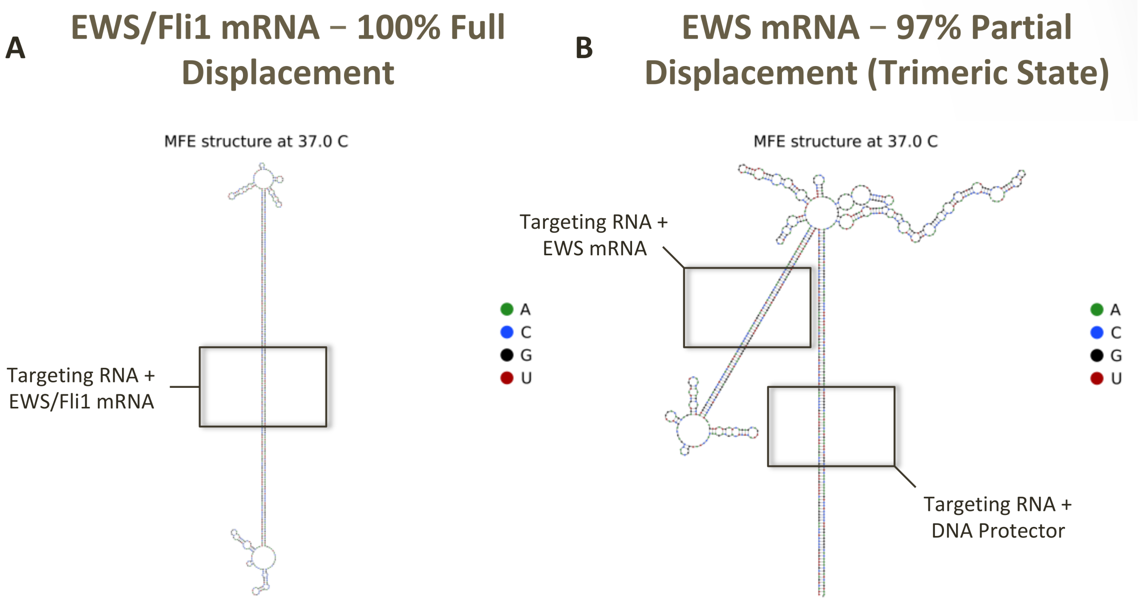


**Figure S6. Trimeric state formation of 200 bp (100/100) EWS/Fli1 targeting RNA.** Secondary structure assessment using NUPACK software reveals [A] complete displacement of the protected RNA in the presence of the EWS/Fli1 mRNA target but [B] only partial displacement and trimer formation in the presence of the EWS or Fli1 mRNA wildtype sequences. The trimer complex represented here depicts targeting RNA, its DNA protector, and EWS mRNA. It should be noted that most secondary structure simulation softwares, including NUPACK, are currently incapable of simulating RNA/DNA binding. Accordingly, the ΔG of any RNA-DNA duplexes were first derived in MATLAB using thermodynamic parameters from the 1995 paper by Sugimoto, et al. The DNA protector was then converted into its equivalent RNA strand for secondary structure derivation via NUPACK, but was truncated (truncation was tested from both the 5' and 3' ends but is depicted from the 3' end for this figure) until the ΔG of the targeting RNA-RNA protector duplex was equivalent to the ΔG of the targeting RNA-DNA protector duplex derived via MATLAB. Arrowheads signify 3' ends.

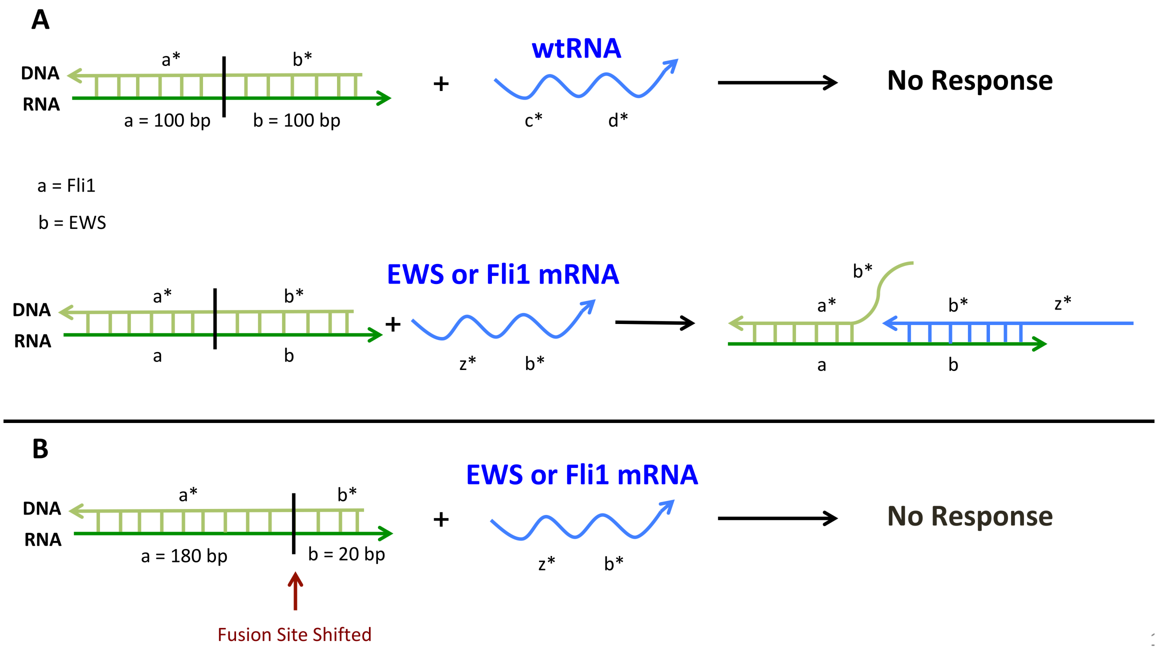


**Figure S7. Targeting strand modification to prevent trimer formation.** To prevent trimer formation, where targeting RNA, its DNA protector, and the target mRNA are all bound, the fusion-site can be shifted away from the [A] center (100/100 bp) and towards the [B] 3' end of the EWS/Fli1 targeting RNA (180/20 bp), kinetically locking the targeting RNA strand and preventing non-specific displacement and activation. Arrowheads signify 3' ends. Asterisks signify complementarity.

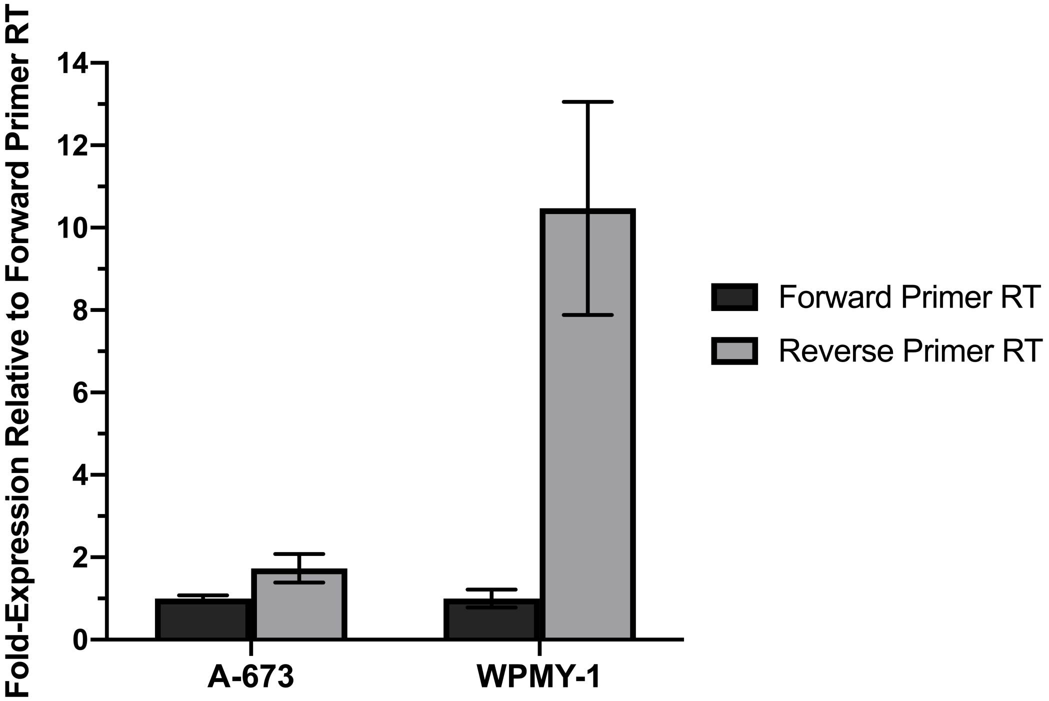


**Figure S8. Utilizing 2-step RT-qPCR to validate the presence of transcribed 2'-U DNA.** A modified 2-step RT-qPCR protocol was used to verify the unique RNA signal observed in Figure 5c. Briefly, during the reverse transcriptase step, either forward or reverse primer alone was added but not both. Following reverse transcription, the reverse transcriptase was heat inactivated and qPCR was performed by adding the missing primer component. The presence of a stronger signal with the use of a reverse primer versus forward primer during cDNA synthesis indicates that the amplified product detected in Figure 5c is the transcribed 2'-U DNA product and not Fli1 mRNA. Error bars represent the standard deviation of replicate conditions.

**Table S1. List of all sequences and primers.**
