## Supplementary Table S1 for "Development of a Novel Class of Self-Assembling dsRNA Cancer Therapeutics: a Proof of Concept Investigation"

| Targeting RNA Strands | Sequence | Notes |
|---|---|---|
| **Scrambled 200 bp** | CAGGCAAUUAAGGGGAACGUAAGUAUAGCGCAAAAAAACAGAGAAAUAGGCGAAUGAAUCUAUUCUUACUGUAUCGAAGAAUGGCCUCGCGGAGGCAUGUGUCA<br>UGCUAGCGUGCGGGGUACUCUAGUUAUCCAUAUGGUCCACAGGACACUCGUUGCUUUCGGAUUUGCCCUUUAUGCGCCGGUUUUCAGCCACGCUUA | |
| **EWS/Fli1 10/10 bp** | UAAGAAGGGUUCUGCUGCCC | |
| **EWS/Fli1 20/20 bp** | GACUGAGUCAUAAGAAGGGUUCUGCUGCCCGUAGCUGCUG | |
| **EWS/Fli1 35/35 bp** | CCAAGCUCCUCUUCUGACUGAGUCAUAAGAAGGGUUCUGCUGCCCGUAGCUGCUGCUCUGUUGGCUAUAU | |
| **EWS/Fli1 60/60 bp** | UGAGGCCAGAAUUCAUGUUAUUGCCCCAAGCUCCUCUUCUGACUGAGUCAUAAGAAGGGUUCUGCUGCCCGUAGCUGCUGCUCUGUUGGCUAUAUUGACUUGGA<br>GCUUGGCUGUAGGAUC | |
| **EWS/Fli1 70/70 bp** | CAGGCAAGGACUUUUGUUGAGGCCAGAAUUCAUGUUAUUGCCCCAAGCUCCUCUUCUGACUGAGUCAUAAGAAGGGUUCUGCUGCCCGUAGCUGCUGCUCUGUU<br>GGCUAUAUUGACUUGGAGCUUGGCUGUAGGAUCCAGUUUGGGG | |
| **EWS/Fli1 100/100 bp** | UUACUGAUCGUUUGUGCCCCUCCAAGGGGAGGACUUUUGUUGAGGCCAGAAUUCAUGUUAUUGCCCCAAGCUCCUCUUCUGACUGAGUCAUAAGAAGGGUUCUG<br>CUGCCCGUAGCUGCUGCUCUGUUGGCUAUAUUGACUUGGAGCUUGGCUGUAGGAUCCAGUUUGGGGUGGGUAACUAGUGGGAGGCUGCUGCCCAUA | |
| **EWS/Fli1 200/190 bp** | UUGCCACAGCUGGAUCUGCCCGCUUCCAGGGUUGGCUAGGCGACUGCUGGUCGGGCCCAGGAUCUGAUACGGAUCUGGCUGGGGCCGUUGCUCUGUAUUCUUAC<br>UGAUCGUUUGUGCCCCUCCAAGGGGAGGACUUUUGUUGAGGCCAGAAUUCAUGUUAUUGCCCCAAGCUCCUCUUCUGACUGAGUCAUAAGAAGGGUUCUGCUGC<br>CCGUAGCUGCUGCUCUGUUGGCUAUAUUGACUUGGAGCUUGGCUGUAGGAUCCAGUUUGGGGUGGGUAACUAGUGGGAGGCUGCUGCCCAUAGCUGCUUUGUU<br>GACCAUAGCUACUCUGCUGUCCAUAGCUGCUCGGUUGCCCAUAGGUGUUCUGCUGAGAGUAACUGCUCUGAUCAUAACU | |
| **EWS/Fli1 180/20 bp Fusion-Shifted** | GGCAACGCUUCCAGGGUUGGCUAGGCGACUGCUGGUCGGGCCCAGGAUCUGAUACGGAUCUGGCUGGGGCCGUUGCUCUGUAUUCUUACUGAUCGUUUGUGCCC<br>CUCCAAGGGGAGGACUUUUGUUGAGGCCAGAAUUCAUGUUAUUGCCCCAAGCUCCUCUUCUGACUGAGUCAUAAGAAGGGUUCUGCUGCCCGUAGCUGCUG | |
| **EWS/Fli1 2'-F 180/20 bp Fusion-Shifted** | 2FG2FGC2FA2FAC2FGCUUCC2FA2FG2FG2FGUU2FG2FGCU2FA2FG2FGC2FG2FACU2FGCU2FG2FGUC2FG2FG2FGCCC2FA2FG2FG2FAUCU2FG2FAU2FAC2FG<br>2FG2FAUCU2FG2FGCU2FG2FG2FG2FGCC2FGUU2FGCUCU2FGU2FAUUCUU2FACU2FG2FAUC2FGUUU2FGU2FGCCCCUCC2FA2FA2FG2FG2FG2FG2FA2FG2F<br>G2FACUUUU2FGUU2FG2FA2FG2FGCC2FA2FG2FA2FAUUC2FAU2FGUU2FAUU2FGCCCC2FA2FA2FGCUCCUCUUCU2FG2FACU2FG2FA2FGUC2FAU2FA2FA2F<br>G2FA2FA2FG2FG2FGUUCU2FGCU2FGCCC2FGU2FA2FGCU2FGCU2FG | 2F = 2'-F |
| **EWS/Fli1 15/15 bp End-Blocked Pilot** | dA*dG*dT*dCdAUAAGAAGGGUUCUGCUGCCCdGdTdA*dG*dC* | * = PS-bond, d = DNA base |
| **EWS/Fli1 100/20 bp Fusion-Shifted End-Blocked** | dA*dA*dT*dGdAGAUCGUUUGUGCCCCUCCAAGGGGAGGACUUUUGUUGAGGCCAGAAUUCAUGUUAUUGCCCCAAGCUCCUCUUCUGACUGAGUCAUAAGAAG<br>GGUUCUGCUGCCCGUAGC*dT*dG*dC*dT*dG | |
| **DNA Protector Strands** | | |
| **Scrambled 200 bp** | TGGCTGAAAACCGGCGCATAAAGGGCAAATCCGAAAGCAACGAGTGTCCTGTGGACCATATGGATAACTAGAGTACCCCGCACGCTAGCATGACACATGCCTCCGCGAGG<br>CCATTCTTCGATACAGTAAGAATAGATTCATTCGCCTATTTCTCTGTTTTTTTGCGCTATACTTACGTTCCCCTTAATTGCCTG | |
| **Scrambled 200 bp 2'-U** | /IDEOXYU/GGC/IDEOXYU/GAAAACCGGCGCA/IDEOXYU/AAAGGGCAAA/IDEOXYU/CCGAAAGCAACGAG/IDEOXYU/G/IDEOXYU/CC/IDEOXYU/G/IDEOXYU/GG<br>ACCA/IDEOXYU/A/IDEOXYU/GGA/IDEOXYU/AAC/IDEOXYU/AGAG/IDEOXYU/ACCCCGCACGC/IDEOXYU/AGCA/IDEOXYU/GACACA/IDEOXYU/GCC/IDEOXYU/<br>CCGCGAGGCCA/IDEOXYU//IDEOXYU/C/IDEOXYU//IDEOXYU/CGA/IDEOXYU/ACAG/IDEOXYU/AAGAA/IDEOXYU/AGA/IDEOXYU//IDEOXYU/CA/IDEOXYU//ID<br>EOXYU/CGCC/IDEOXYU/A/IDEOXYU//IDEOXYU//IDEOXYU/C/IDEOXYU/C/IDEOXYU/G/IDEOXYU//IDEOXYU//IDEOXYU//IDEOXYU//IDEOXYU//IDEOXYU//IDE<br>OXYU/GCGC/IDEOXYU/A/IDEOXYU/AC/IDEOXYU//IDEOXYU/ACG/IDEOXYU//IDEOXYU/CCCC/IDEOXYU//IDEOXYU/AA/IDEOXYU//IDEOXYU/GCC/IDEOXYU/<br>G | /IDEOXYU/ = 2'-U |
| **Scrambled 200 bp 2'-O Methyl** | TGGCT2MGAAAACCGGC2MGCATAAAG2MGGCAAATCC2MGAAAGCAAC2MGAGTGTCCT2MGTGGACCATATG2MGATAACTAGA2MGTACCCCGCAC2MGCTAGC<br>ATGACACAT2MGCCTCC2MGCGAGGCCATTCTTC2MGATACA2MGTAAGAATA2MGATTCATTC2MGCCTATTTCTCT2MGTTTTTTGC2MGCTATACTTAC2MGTTCCC<br>CTTAATT2MGCCT2MG | 2MG = 2'-O Methyl |
| **EWS/Fli1 100/100 bp** | AGCCTCCCACTAGTTACCCACCCCAAACTGGATCCTACAGCCAAGCTCCAAGTCAATATAGCCAACAGAGCAGCAGCTACGGGCAGCAGAACCCTTCTTATGACTCAGTCAG<br>AAGAGGAGCTTGGGGCAATAACATGAATTCTGGCCTCAACAAAAGTCCTCCCCTTGGAGGGGCACAAACGATCAGTAA | |
| **EWS/Fli1 180/20 bp Fusion-Shifted** | GGGCAGCAGAACCCTTCTTATGACTCAGTCAGAAGAGGAGCTTGGGGCAATAACATGAATTCTGGCCTCAACAAAAGTCCTCCCCTTGGAGGGGCACAAACGATCAGTAAG<br>AATACAGAGCAACGGCCCCAGCCAGATCCGTATCAGATCCTGGGCCCGACCAGCAGTCGCCTAGCCAACCCTGGAAGCGTTGCCTGCCC | |
| **EWS/Fli1 15/15 bp End-Blocked Pilot** | G*G*G* C*AG CAG AAC CC/IDEOXYU/ /IDEOXYU/C/IDEOXYU/ /IDEOXYU/A/IDEOXYU/ GAC /IDEOXYU/A | /IDEOXYU/ = 2'-U |
| **EWS/Fli1 100/20 bp Fusion-Shifted End-Blocked** | ACG GGC AGC AGA ACC C/IDEOXYU//IDEOXYU/ C/IDEOXYU//IDEOXYU/ A/IDEOXYU/G AC/IDEOXYU/ CAG /IDEOXYU/CA GAA GAG GAG<br>C/IDEOXYU//IDEOXYU/ GGG GCA A/IDEOXYU/A ACA /IDEOXYU/GA A/IDEOXYU//IDEOXYU/ C/IDEOXYU/G GCC /IDEOXYU/CA ACA AAA G/IDEOXYU/C<br>C/IDEOXYU/C CCC /IDEOXYU//IDEOXYU/G GAG GGG CAC AAA CGA /IDEOXYU/C/IDEOXYU/ CA/IDEOXYU/ /IDEOXYU/ | /IDEOXYU/ = 2'-U |
| **RNA Protector Strands** | | |
| **Scrambled 200 bp s2U** | S2UGGCS2UGAAAACCGGCGCAS2UAAAGGGCAAAS2UCCGAAAGCAACGAGS2UGS2UCCS2UGS2UGGACCAS2UAS2UGGAS2UAACS2UAGAGS2UACCCCGCACG<br>CS2UAGCAS2UGACACAS2UGCCS2UCCGCGAGGCCAS2US2UCS2US2UCGAS2UACAGS2UAAGAAS2UAGAS2US2UCAS2US2UCGCCS2UAS2US2US2UCS2UCS2U<br>GS2US2US2US2US2US2US2UGCGCS2UAS2UACS2US2UACGS2US2UCCCCS2US2UAAS2US2UGCCS2UG | S2U = 2-Thiouridine |
| **Scrambled 200 bp s4U GU Wobble** | S4UGGCS4UGAGAACCGGCGCGS4UAAAGGGCGAAS4UCCGAGAGCAACGGGS4UGS4UCCS4UGS4UGGACCGS4UAS4UGGAS4UAGCS4UAGAGS4UGCCCCGCACG<br>CS4UGGCAS4UGACGCAS4UGCCS4UCCGCGGGGCCGS4US4UCS4US4UCGAS4UGCAGS4UAAGAGS4UAGAS4US4UCAS4US4UCGCCS4UGS4US4US4UCS4UCS4<br>UGS4US4US4US4US4US4US4UGCGCS4UGS4UACS4US4UACGS4US4UCCCCS4US4UGAS4US4UGCCS4UG | S4U = 4-Thiouridine |

| | | |
|---|---|---|
| **Scrambled 200 bp 5-mCTP** | UGG5MCUGAAAA5MC5MCGG5MCG5MCAUAAAGGG5MCAAAU5MC5MCGAAAG5MCAA5MCGAGUGU5MC5MCUGUGGA5MC5MCAUAUGGAUAA5MCUAG<br>AGUA5MC5MC5MC5MCG5MCA5MCG5MCUAG5MCAUGA5MCA5MCAUG5MC5MCU5MC5MCG5MCGAGG5MC5MCAUU5MCUU5MCGAUA5MCAGUAAGA<br>AUAGAUU5MCAUU5MCG5MC5MCUAUUU5MCU5MCUGUUUUUUUG5MCG5MCUAUA5MCUUA5MCGUU5MC5MC5MC5MCUUAAUUG5MC5MCUG | 5MC = 5-methylcytidine |
| **Truncated mRNA Sense Strands** | | |
| **EWS/Fli1 400 bp** | AGTTATGAT CAGAGCAGTT ACTCTCAGCAGAACACCTAT GGGCAACCGA GCAGCTATGG ACAGCAGAGT AGCTATGGTC AACAAAGCAGCTATGGGCAG<br>CAGCCTCCCA CTAGTTACCC ACCCCAAACT GGATCCTACA GCCAAGCTCCAAGTCAATAT AGCCAACAGA GCAGCAGCTA CGGGCAGCAG AACCCTTCT TATGACTCAG<br>TCAGAAGAGGAGCTTGGGGC AATAACATGA ATTCTGGCCT CAACAAAAGT CCTCCCCTTG GAGGGGCACA AACGATCAGT AAGAATACAG AGCAACGGCC<br>CCAGCCAGAT CCGTATCAGA TCCTGGGCCC GACCAGCAGT CGCCTAGCCA ACCCTGGAAG CGGGCAGATC CAGCTGTGGC AA | |
| **EWS 400 bp** | AGTTATGAT CAGAGCAGTT ACTCTCAGCA GAACACCTAT GGGCAACCGA GCAGCTATGG ACAGCAGAGT AGCTATGGTC AACAAAGCAG CTATGGGCAG<br>CAGCCTCCCA CTAGTTACCC ACCCCAAACT GGATCCTACA GCCAAGCTCC AAGTCAATAT AGCCAACAGA GCAGCAGCTA CGGGCAGCAG AGTTCATTCC<br>GACAGGACCA CCCCAGTAGC ATGGGTGTTT ATGGGCAGGA GTCTGGAGGA TTTTCCGGAC CAGGAGAGAA CCGGAGCATG AGTGGCCCTG ATAACCGGGG<br>CAGGGGAAGA GGGGGATTTG ATCGTGGAGG CATGAGCAGA GGTGGGCGGG GAGGAGGACG CGGTGGAATG GGCGCTGGAG AGCGAGGTGG CTTCAATAA | |
| **Fli1 400 bp** | GGAGATCG ACACATCCTT TTTCCAGAAC ATGGATGGCA AGGAACTGTG TAAAATGAAC AAGGAGGACT TCCTCCGCGC CACCACCCTC TACAACACGG<br>AAGTGCTGTT GTCACACCTC AGTTACCTCA GGGAAAGTTC ACTGCTGGCC TATAATACAA CCTCCCACAC CGACCAATCC TCACGATTGA GTGTCAAAGA<br>AGACCCTTCT TATGACTCAG TCAGAAGAGG AGCTTGGGGC AATAACATGA ATTCTGGCCT CAACAAAAGT CCTCCCCTTG GAGGGGCACA AACGATCAGT<br>AAGAATACAG AGCAACGGCC CCAGCCAGAT CCGTATCAGA TCCTGGGCCC GACCAGCAGT CGCCTAGCCA ACCCTGGAAG CGGGCAGATC CAGCTGTGGC AA | |
| **Primers** | | |
| **EWS/Fli1 mRNA** | F primer: CAGCAGAGTAGCTATGGTCAAC; R primer: GGCCGTTGCTCTGTATTCTTA |  |
| **EWS mRNA** | F primer: CAGCAGAGTAGCTATGGTCAAC; R primer: TCCGGAAAATCCTCCAGAC | |
| **Fli1 mRNA** | F primer: GGATGGCAAGGAACTGTGTA; R primer: GGCCGTTGCTCTGTATTCTTA | |
| **GAPDH mRNA** | F primer: GGGTGTGAACCATGAGAAGTAT; R primer: AGTAGAGGCAGGGATGATGT | |
| **2'-U Transcribed DNA RT-qPCR** | F primer: GAGGACTTTTGTTGAGGCCAG; R primer = CAGTCAGAAGAGGAGCTTGGG | |
| **PKR mRNA** | F primer: GGTACAGGTTCTACTAAACAGG; R primer: GAAAACTTGGCCAAATCCACC | |
| **RIG-I mRNA** | F primer: TGTTCTCAGATCCCTTGGATG; R primer: CACTGCTCACCAGATTGCAT | |
| **OAS3 mRNA** | F primer: CCGAACTGTCCTGGGCCTGATCC; R primer: CCCATTCCCCAGGTCCCATGTGG | |
